# Late-Stage Posttranslational Assembly of Fosfazinomycins

**DOI:** 10.64898/2026.09.23.753898

**Authors:** Lide Cha, Zedu Huang, Chandrashekhar Padhi, Kwo-Kwang A. Wang, Wilfred A. van der Donk

## Abstract

The fosfazinomycins are phosphonate natural products with antifungal activity. Their main scaffold is composed of a phosphonate moiety attached to the carboxylate of arginine via a hydrazine linkage. Previous studies have elucidated a convergent biosynthetic pathway that independently assembles phosphonate and hydrazine synthons, but how these building blocks are then connected could not be determined. In this work, we provide the final missing steps in fosfazinomycin biosynthesis by revealing an unexpected engagement of biosynthetic machinery that is typically involved in ribosomally synthesized and post-translationally modified peptides (RiPPs). An asparagine synthetase-like (AS-like) enzyme catalyzes the installation of hydrazine onto the carboxylate of Arg at the C-terminus of a short ribosomally synthesized precursor peptide. The terminal nitrogen of the resulting peptide hydrazide is methylated, and the phosphonate moiety is activated to a triphosphate-like intermediate through two separate kinase catalyzed phosphorylation steps. A nucleotidyl transferase then catalyzes the ligation of the two fragments to generate the mature fosfazinomycin scaffold on a peptide. Aminopeptidase cleavage of this peptide then yields fosfazinomycin B (fosB), which serves as substrate for a valinyl-tRNA dependent reaction to afford fosfazinomycin A (fosA). This work demonstrates an unprecedented example of the convergence of RiPP and phosphonate biosynthetic logic, an enzymatic route to peptide hydrazides that are widely used in peptide ligation chemistry, and an unusual activation sequence for conjugation of phosphonates.

## Main

Phosphonates are a commercially significant class of compounds defined by the presence of a carbon–phosphorus (C–P) bond^1^. They exhibit diverse bioactivities including antibacterial, herbicidal, antifungal and antiparasitic activities^2,3^, such as the broad-spectrum herbicide glufosinate^4^ and the broad-spectrum antibiotic fosfomycin^5^. The bioactivities of these molecules typically result from the structural resemblance of the phosphonate moiety to carboxylate or phosphate groups of primary metabolites^1^. As such, they competitively inhibit enzymes that act on these metabolites^6^. Over 100 phosphonate natural products have been identified to date^3^. Almost all of the currently characterized phosphonate biosynthetic pathways initiate with the conversion of phosphoenolpyruvate (PEP) to 3-phosphonopyruvate (PnPy) catalyzed by PEP mutase (PepM)^7,8^. Following the C–P bond formation step, the pathways branch and include transamination, decarboxylation, reduction and condensation reactions (Supplementary Fig. 1)^3^.

A growing subgroup of phosphonate natural products is the phosphonopeptides (Fig. 1A)^3^. These compounds are typically composed of a phosphonate scaffold that is attached to a short peptide. Multiple studies have demonstrated that phosphonopeptides function as “Trojan Horse” molecules where appending the phosphonate to a peptide promotes uptake by target organisms. Upon import by peptide permeases, the bioactive phosphonate warhead is released by cellular peptidases^9–11^. The biosynthetic machinery involved in assembling these peptide or pseudopeptide scaffolds have been studied for a few phosphonopeptides, with the peptide bonds formed by ATP-grasp enzymes^12–14^, aminoacyl-tRNA-dependent ligases^15–18^ or nonribosomal peptide synthetases^19^.

**Figure 1.**
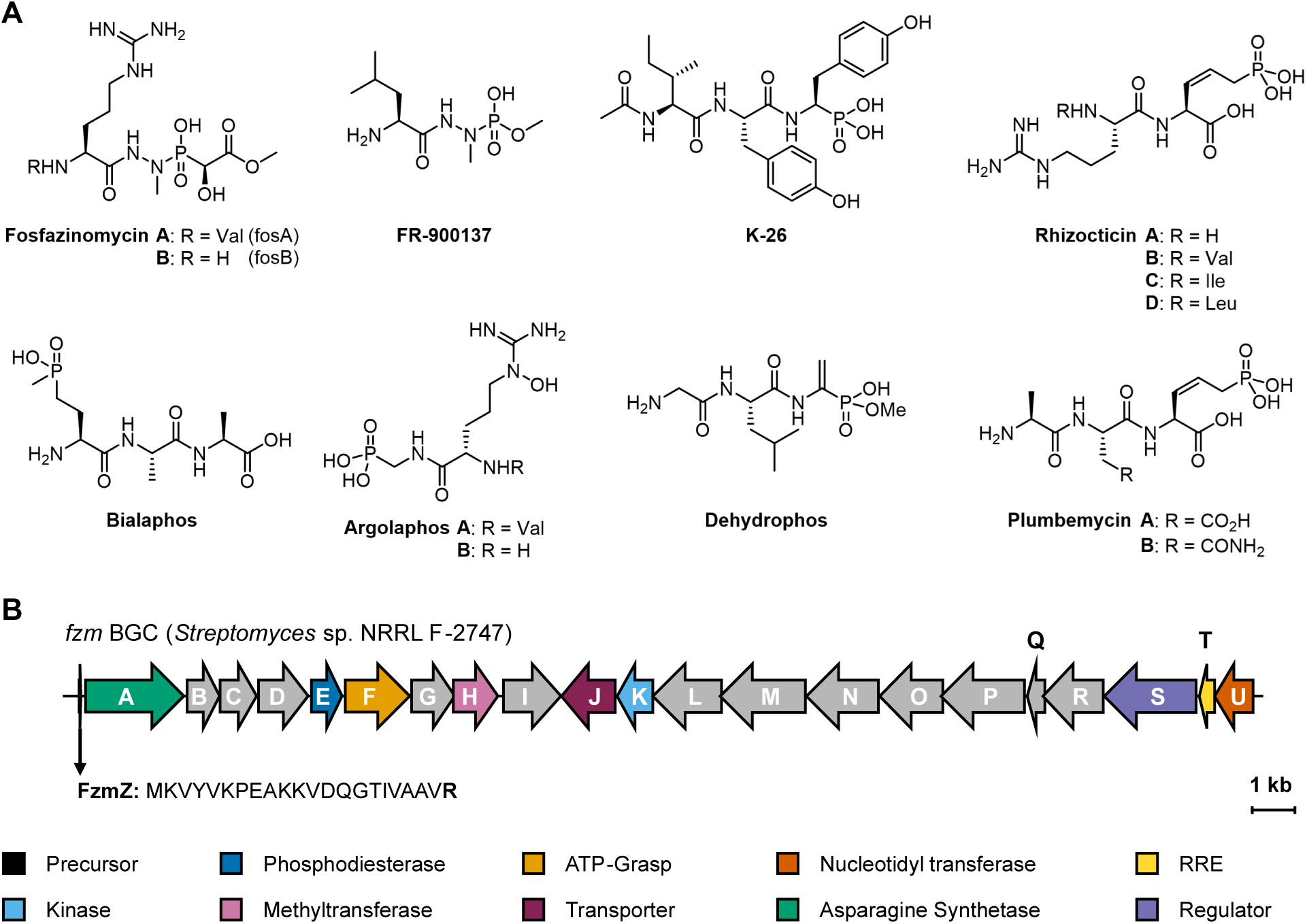
Phosphonopeptides and the reannotated fosfazinomycin BGC. **(A)** The chemical structures of representative phosphonopeptide natural products and the fosfazinomycin analog FR-900137. **(B)** The reannotated fosfazinomycin BGC from *Streptomyces* sp. NRRL F-2747. The amino acid sequence of the precursor peptide FzmZ is shown highlighting the C-terminal arginine residue. Genes colored in gray have been characterized in previous studies^16,21–24^.

The antifungal fosfazinomycin is a unique member of the phosphonopeptides^20^. A hydrazine moiety connects the carboxylate of arginine and the phosphonic acid of the phosphonate (Fig. 1A). Previous studies on the enzymes encoded in the fosfazinomycin biosynthetic gene cluster (BGC) have established how the phosphonate moiety as well as the hydrazine fragment are biosynthesized^16,21–24^. However, how these fragments are connected to yield fosfazinomycins has remained elusive.

In this work, we present the missing steps for the biosynthesis of fosfazinomycin. These results surprisingly demonstrate that the assembly of the fosfazinomycin backbone uses biosynthetic machinery of ribosomally synthesized and post-translationally modified peptides (RiPPs)^25^. The asparagine synthetase-like (AS-like) enzyme FzmA transfers hydrazine from glutamylhydrazine to the C-terminal Arg residue of a short ribosomally synthesized precursor peptide FzmZ that is encoded in the BGC, but that because of its short length (22 amino acids) had escaped identification in previous studies. Following FzmH-catalyzed *N*-methylation of the peptidyl hydrazide, the ligation of the phosphonate is facilitated by the combined effort of FzmFKU using an unprecedented activation mechanism. The two kinases FzmF and FzmK catalyze the sequential phosphorylation of methyl 2-hydroxy-2-phosphono-acetate (Me-HPnA) to form a phosphonate diphosphate. The resulting activated phosphonate is then ligated to the peptide hydrazide by the nucleotidyl transferase FzmU. A housekeeping aminopeptidase in the producing organism is likely responsible for the proteolytic liberation of fosfazinomycin B, which was previously shown to be the substrate of FzmI to generate fosfazinomycin A^22^. This remarkably complex pathway appears to have evolved to never generate the phosphonohydrazide as an intermediate, suggesting that it may be the cytotoxic compound, in line with other Trojan horse antibiotics^9,15,26–29^. Overall, these results highlight the versatility of RiPP biosynthetic machinery in the production of natural products beyond canonical peptides, provide an enzymatic route to peptide hydrazides that are widely used in synthetic peptide chemistry, and provide new insights into the biosynthesis of phosphonate natural products. Our results suggest that the clinically used antibiotic fosfomycin may undergo similar conjugation chemistry in its producing organism.

## Results

### Bioinformatic Identification and Reannotation of the *fzm* BGC

During a recent bioinformatic investigation of AS-like enzymes related to RiPP biosynthesis, we generated a sequence similarity network (SSN) (Extended Data Fig.1)^30^. The SSN contains a cluster of enzymes (cluster 58) that possess a fused C-terminal RiPP precursor recognition element (RRE)^31^. Genome neighborhood analysis^32^ of the AS-like enzymes from this cluster revealed that they often co-occur with discrete RREs (PF05402, 79%), nucleotidyl transferases (IPR043519, 62%), AurF-like dioxygenases (PF11583, 60%) and MFS transporter (PF07690, 54%) (Extended Data Fig. 2A). Initial attempts to identify potential precursor peptides that might bind to the RREs by retrieving short open reading frames (ORFs) ranging from 30 to 150 amino acids in the genome neighborhood using the RODEO webtool^33^ were unsuccessful, but careful manual examination led to the identification of a 21/22-mer precursor peptide in each candidate BGC. The peptides were of interest because they contain a conserved N-terminal YxxP motif that is often used for RRE engagement^34,35^ (Extended Data Fig. 2B, Supplementary Fig. 2A).

FzmA, an AS-like enzyme that was previously identified in the biosynthetic pathway of fosfazinomycins, was unexpectedly present in this enzyme cluster in the SSN (Extended Data Fig. 1). A previous study proposed that FzmA is responsible for transferring hydrazine from glutamylhydrazide to the carboxylate of arginine to construct argininylhydrazide. While the hydrazide formation activity of FzmA was not observed when this hypothesis was tested, FzmA was shown to liberate hydrazine from glutamylhydrazide^23^. The presence of an RRE in FzmA and the observation that the abovementioned peptide encoded near it contains a C-terminal Arg residue (Fig. 1B) led us to suspect that the previous unsuccessful attempts to reconstitute FzmA activity was the result of using the incorrect substrate. We constructed an AlphaFold3^36^ structural model of FzmA from *Streptomyces* sp. NRRL F-2747, which agrees with the presence of a fused RRE domain at the C-terminus of the protein (Extended Data Fig. 2C). The 22-amino acid ORF adjacent to *fzmA,* the hypothetical precursor peptide, contains the conserved N-terminal YxxP motif and a C-terminal VR motif, which is consistent with the amino acid compositions of fosfazinomycin A and B (Fig. 1A). Inspection of all orthologous *fzm* BGCs unveiled that such a precursor peptide is encoded in all such BGCs (Extended Data Fig. 2D, Supplementary Fig. 2B). Together, these findings strongly suggested that the short 22-mer ribosomal peptide (which we here denote FzmZ) is the actual substrate of FzmA.

The suggested connection of the *fzm* BGC to RiPP biosynthesis also led us to examine the genome neighborhood of the BGC for other RiPP elements that may have gone unnoticed during prior studies. A gene pair encoding a discrete RRE (*fzmT*) as well as a hypothetical nucleotidyl transferase (*fzmU*), was identified immediately outside of the previously defined boundary of the *fzm* BGC in *Streptomyces* sp. NRRL F-2747 (Fig. 1B). This gene pair is also found adjacent to the *fzm* BGCs from other known fosfazinomycin producing strains as well as in low-identity homologous *fzm* BGCs (Supplementary Fig. 3).

### FzmA and FzmH generate a Peptidyl Methyl Hydrazide

We first tested whether FzmZ is a substrate of the hypothetical hydrazide synthetase FzmA. Original attempts to heterologously express *N*-terminally His_6_-tagged FzmZ in *Escherichia coli* were not successful. We therefore prepared FzmZ by solid phase peptide synthesis. Incubation of purified FzmA with FzmZ, ATP, Mg^2+^ and glutamylhydrazine, and subsequent LC-HRMS analysis revealed the formation of a new species with a mass shift of +14 Da (Fig. 2). The product peptide exhibited the same retention time and fragmentation pattern as a synthetic FzmZ standard with a C-terminal hydrazide (denoted FzmZ-NHNH_2_, Fig. 3A, Supplementary Fig. 4, 5). Together, these observations suggest that FzmA catalyzes the conjugation of hydrazine to the C-terminus of a peptide substrate. A brief substrate scope analysis of FzmA showed that the enzyme can also accept L-Gln and L-Glu-NHOH as co-substrates to catalyze amidation and hydroxylamidation of the C-terminus of FzmZ, respectively (Extended Data Fig. 3).

**Figure 2.**
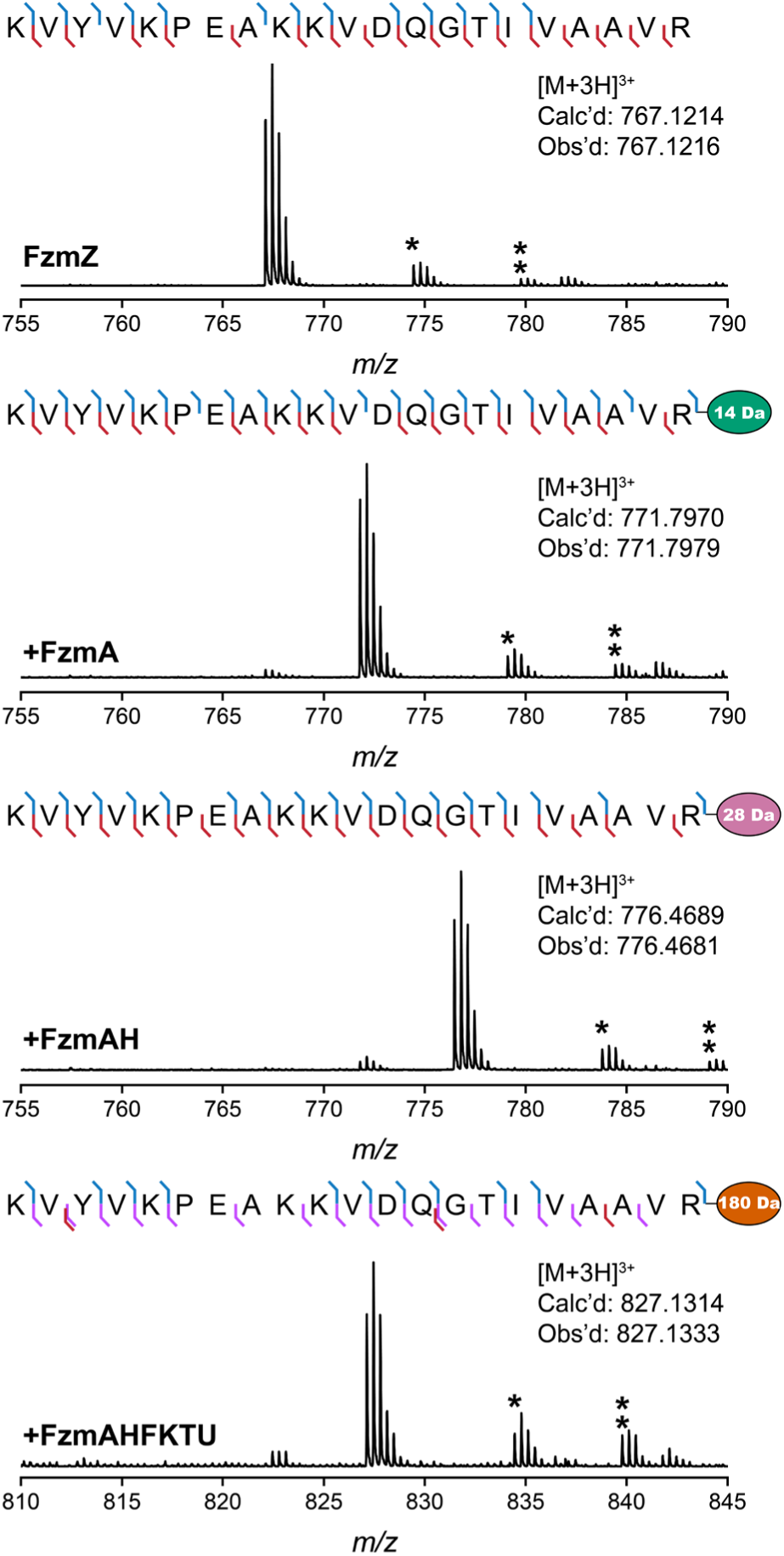
HR-MS/MS analysis of unmodified precursor peptide FzmZ and enzyme-modified FzmZ. The MS/MS fragmentation pattern of each peptide is shown (see Supplementary Fig. 4-5 for MS/MS spectra). Annotation of the tandem mass spectrum was performed with the interactive peptide spectral annotator^37^. The modifications introduced to the C-terminal carboxylate of the Arg residue are labelled as colored ellipses with the masses labelled. Asterisk denotes [M+2H+Na]^3+^, double asterisk denotes [M+2H+K]^3+^.

**Figure 3.**
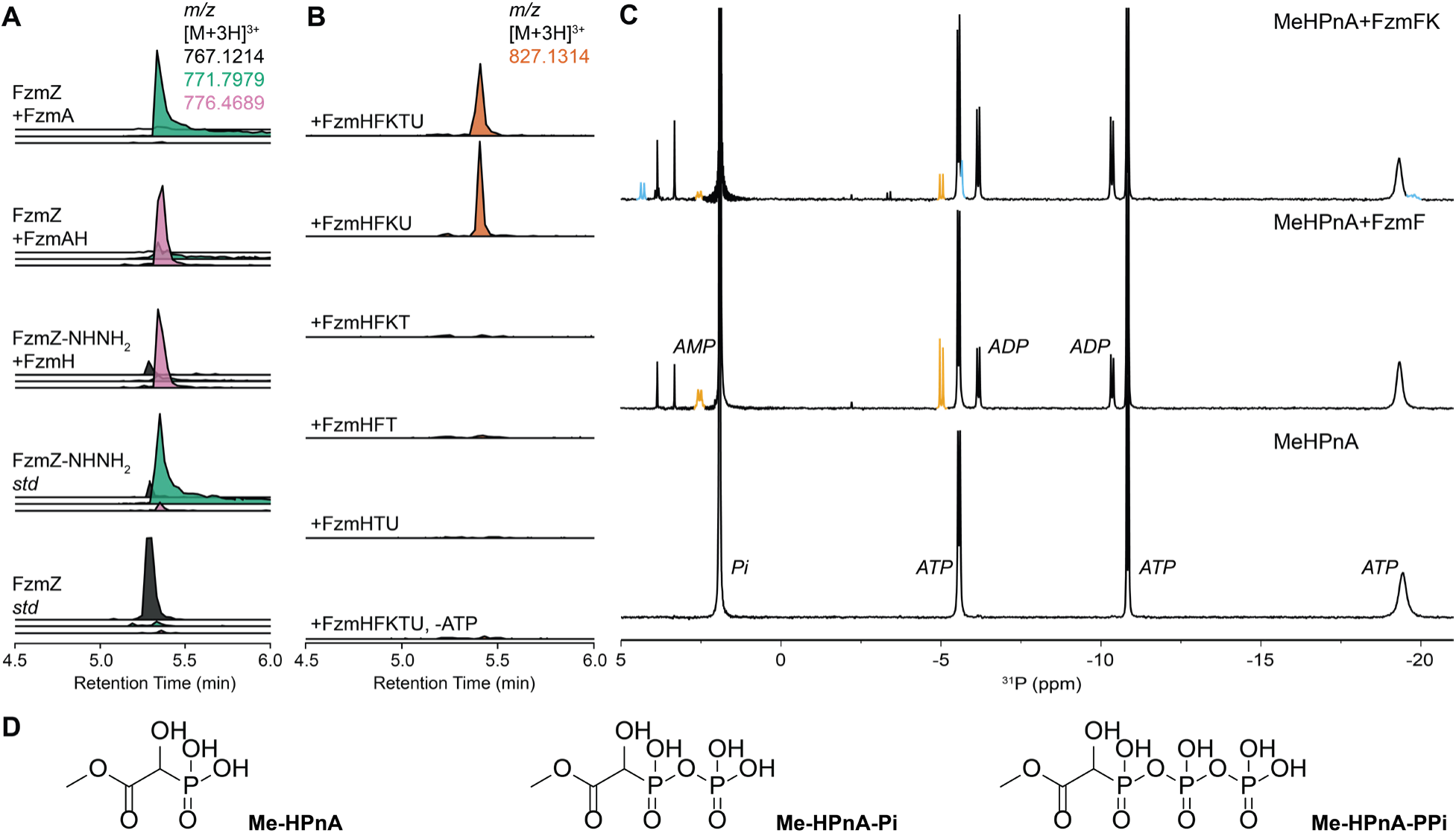
Functional validation of FzmA, FzmH, FzmF, FzmK, FzmT and FzmU. **(A)** LC-HRMS analysis of *in vitro* reactions of FzmA and FzmH with synthetic FzmZ or FzmZ-NHNH_2_ (calculated mass for FzmZ [M+3H]^3+^: 767.1214, FzmZ-NHNH_2_ [M+3H]^3+^: 771.7970, FzmZ-NHNHMe [M+3H]^3+^: 776.4689). See Fig. 2 for HRMS spectra. Glutamylhydrazide and ATP were included in assays with FzmA or FzmH, respectively. **(B)** LC-HRMS analysis of the *in vitro* reaction of FzmFKTU with FzmZ-NHNHMe, Me-HPnA, and ATP (calculated mass for FzmZ-fosB [M+3H]^3+^: 827.1314). **(C)** ^31^P NMR analysis of the *in vitro* reactions catalyzed by FzmF and FzmK with Me-HPnA in the presence of ATP. The spectrum is zoomed in to the –22 to 5 ppm window and as such, the peak representing Me-HPnA (∼9 ppm) is not shown, see Extended Fig. 4 and Supplementary Fig. 9 for spiking experiments with standards. The peaks of Me-HPnA-Pi are highlighted in yellow and the peaks of Me-HPnA-PPi are highlighted in blue. **(D)** Structures of Me-HPnA, Me-HPnA-Pi and Me-HPnA-PPi.

FzmH, an *S*-adenosylmethionine (SAM) dependent methyltransferase, was previously shown to methylate the terminal nitrogen of argininylhydrazide, but not desmethyl-fosfazinomycin B (Supplementary Fig. 6)^16^. We therefore tested whether the argininylhydrazide at the C-terminus of FzmZ would also be a substrate of FzmH. FzmH and SAM were added to the FzmA-FzmZ assay, resulting in the emergence of a new species with a +14 Da mass shift from the FzmZ-NHNH_2_ peptide (Fig. 2, 3A). The same reaction outcome was also observed when synthetic FzmZ-NHNH_2_ was incubated with FzmH and SAM (Fig. 3A). Tandem MS analysis assigned the 14 Da mass gain to the C-terminal hydrazide moiety (Fig. 2, Supplementary Fig. 5). Therefore, FzmH can install a methyl group on the terminal nitrogen of FzmZ-NHNH_2_ (Fig. 5, product denoted FzmZ-NHNHMe).

### Elucidation of the phosphonate ligation mechanism

We next sought to identify the enzymes responsible for constructing the N–P bond in fosfazinomycins. Two enzymes encoded in the cluster, FzmF (ATP-grasp enzyme) and FzmU (nucleotidyl transferase) were prime candidates for this activity, as reported members of these protein families possess ATP-dependent ligase activities^12–14,38^. When different combinations of these enzymes were incubated with the FzmZ-NHNHMe peptide (obtained from the FzmH assay), the ligation of the phosphonate (Me-HPnA, Fig. 3D) onto FzmZ-NHNHMe was observed when FzmF, FzmK, FzmT and FzmU as well as ATP and Mg^2+^ were present in the assay (Fig. 3B, product denoted FzmZ-fosB). Leaving out the stand-alone RRE FzmT did not impact the ligation outcome (Fig. 3B), but all other enzymes were required. The NTP selectivity of the ligation activity was also evaluated, with reduced activity observed with CTP, and significantly diminished activity with GTP and UTP. Therefore, ATP appears to be the preferred co-substrate for the N–P bond formation reaction (Extended Data Fig. 4).

We then investigated the individual activities of the enzymes. Incubation of FzmF with Me-HPnA in the presence of ATP and subsequent ^31^P NMR analysis revealed the emergence of two new doublets with chemical shifts of 2.6 and –5.0 ppm (Fig. 3C). They shared a coupling constant of ∼ 25 Hz, suggesting that these two peaks likely originate from two coupled phosphorus nuclei. The chemical shift and coupling constant are similar to the two-bond coupling system (^2^*J*_PP_) of ADP^39^. Therefore, FzmF was postulated to catalyze the phosphorylation of Me-HPnA on the phosphonic acid moiety to yield Me-HPnA-Pi (Fig. 3D).

Inclusion of FzmK in the assay resulted in a more complex product profile. The FzmF product peak declined in intensity with the concurrent occurrence of a set of peaks with chemical shifts of 4.3, –5.6 and –19.8 ppm (Fig. 3C). This set of peaks are similar in chemical shifts to triphosphate compounds like ATP^39^. Two of the peaks partially overlapped with ATP signals preventing coupling constant analysis. To verify the structure of the new product, we synthetically prepared a standard of pyrophosphorylated Me-HPnA (Me-HPnA-PPi, Fig. 3D, Supplementary Fig. 7,8). Spiking of the synthetic standard into the FzmFK assay confirmed the assignment of the three new peaks to the three phosphorus atoms of Me-HPnA-PPi (Extended Fig. 4). Collectively, these observations demonstrate that FzmF and FzmK are kinases that sequentially phosphorylate Me-HPnA, yielding a triphosphate-like Me-HPnA-PPi (Fig. 3D), and that the nucleotidyltransferase FzmU uses this activated phosphonate to forge the N-P bond to FzmZ-NHNHMe (Figure 4).

**Figure 4.**
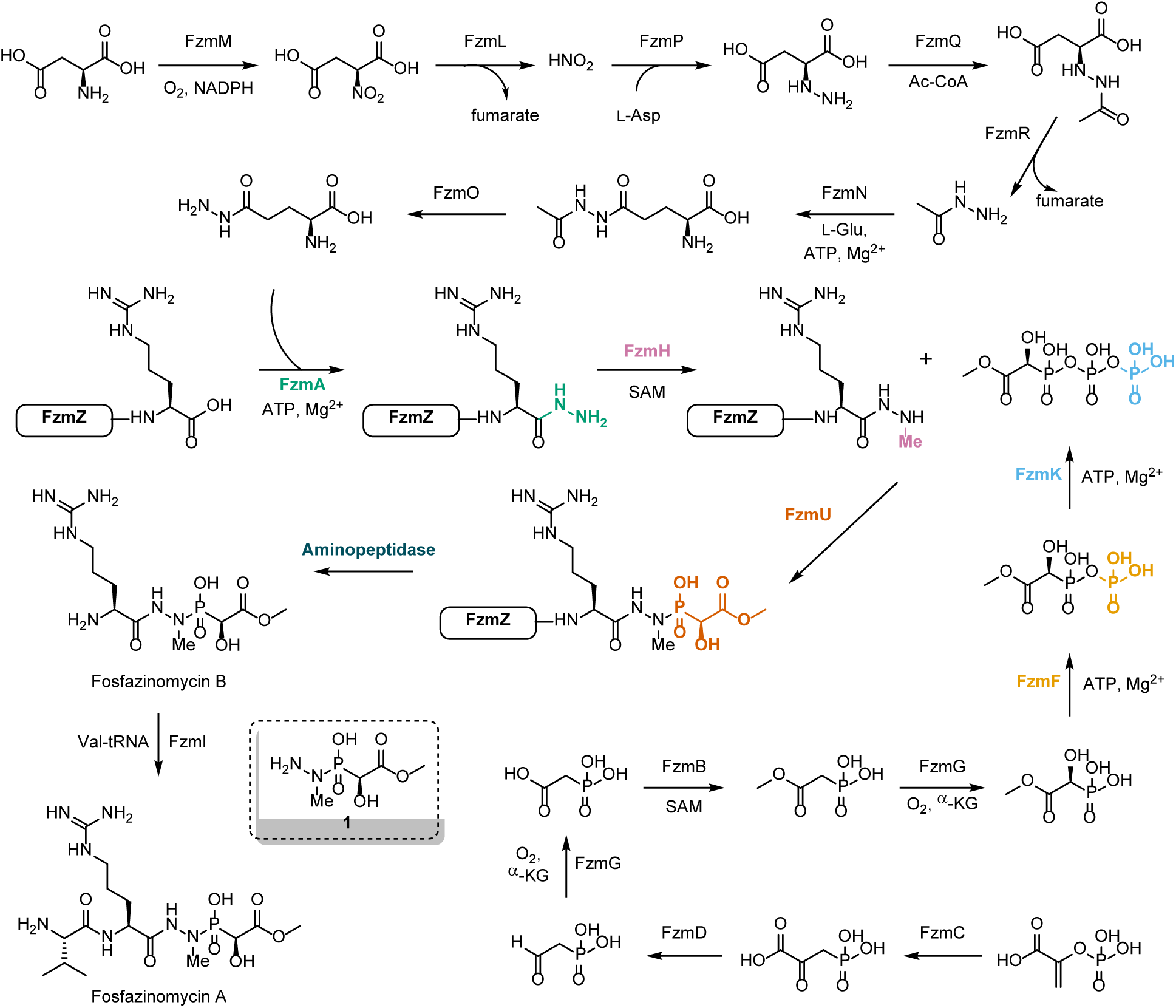
Proposed biosynthetic pathway of fosfazinomycins. Enzymes characterized in this work are highlighted in colored bold text. The structure of the proposed active moiety **1** of fosfazinomycins is shown in the dashed box.

### Attempts to Identify the Proteolysis Mechanism in Fosfazinomycin Biosynthesis

A protease or peptidase that liberates mature fosfazinomycins from the precursor peptide is not encoded in the BGC. We examined the genome of *Streptomyces* sp. NRRL F-2747 for hypothetical protease candidates. An S9 peptidase is encoded 3.1 kilobases upstream of *fzmU*, but it is not conserved. Therefore, this peptidase is unlikely dedicated to the biosynthesis of fosfazinomycins. No other hypothetical peptidases were identified in the 100 kb gene fragment where the *fzm* BGC is located, thus we propose that a general aminopeptidase cleaves the amide bonds until it reaches the C-terminal hydrazide linkage to generate fosfazinomycin B. To test this hypothesis, we treated the product of the FzmFKU-catalyzed conjugation reaction of FzmZ-NHNHMe with Me-HPnA sequentially with endopeptidase LysC and aminopeptidase I from *Streptomyces griseus*. LC-HRMS analysis of the digestion reaction showed the formation of a species with an [M+H]^+^ value of 355.1489. This product shared the same MS/MS fragmentation pattern and retention time with one of the diastereomers of synthetic fosfazinomycin B standard that was prepared with racemic Me-HPnA^40^ (Supplementary Fig. 10). On the other hand, the production of fosfazinomycin A was not detected (Supplementary Fig. 11). This finding is consistent with the previously reported activity of FzmI, which was shown to ligate L-Val to fosfazinomycin B to produce fosfazinomycin A^22^.

## Discussion

The BGC that encodes fosfazinomycin biosynthesis was discovered more than a decade ago^21^ and was surprisingly large for such a small molecule. With this study, we show that the BGC is even larger than previously realized as several enzymes and a scaffolding peptide that were not included in previous studies perform critical functions. With the full biosynthetic route toward fosfazinomycins finally elucidated (Fig. 4)^16,21–24^, a potential explanation can be postulated for the rather unusual pathway. First, and most importantly, the phosphonohydrazide **1** (Fig. 4) that likely is released upon uptake of fosA or fosB by a target organism is never made as a biosynthetic intermediate, thus potentially circumventing toxicity to the producing organism. A recent study on sinefungin biosynthesis provides another example of a seemingly unnecessary long pathway that prevents production of the biologically active material within the cell^41^. The convergent biosynthetic pathway of fosfazinomycin, unique amongst phosphonate natural products that are typically made by linear pathways starting with P–C bond formation, provides the opportunity to link Me-HPnA to Arg-NHNHMe in a manner that prevents production of **1**. This hypothesis also may explain why the compound is assembled on a scaffold peptide FzmZ by repurposing RiPP biosynthetic machinery for the biosynthesis of phosphonate natural products. The requirement for three RiPP-related ORFs explains why in previous attempts to heterologously reconstitute the pathway in *Streptomyces lividans,* only the production of Me-HPnA was observed^21^. Our findings also provide insights into how the leucyl-methylhydrazide moiety of FR-900137^42^ (Fig. 1A) is likely assembled.

The recruitment of an asparagine synthetase-like enzyme and a new activation mechanism for phosphonate chemistry on phosphorus are the key unique steps resulting in fosfazinomycin biosynthesis. AS-like enzymes are involved in amidation^43^ and lactamization^44,45^ of natural products, and more recently they were demonstrated to transfer hydroxylamine to the carboxylate of polyketides^46^ and install a nitrile group on the C-terminus of a ribosomal peptide^30^. The ability of FzmA to catalyze both hydrazide and hydroxyamide formation at the C-terminus of a peptide suggests it may have value in biocatalysis. Peptide C-terminal hydrazides have found extensive use as intermediates for peptide native chemical ligations^47–49^, whereas peptidyl hydroxyamides are widely used as inhibitors of metalloproteases^50^. Investigation of the substrate tolerance of FzmA may unveil its potential as a biocatalyst. We note that peptide hydrazides have also been made enzymatically with sortases^51^ and asparaginyl endopeptidases^52^.

This study also introduces a new mechanism of biosynthetic elaboration of phosphonate structures that borrows from nucleotide chemistry. In natural product biosynthesis, phosphorylation is a prevalent strategy for substrate activation, which enables downstream ligation^53^ and dehydration^25^ activities. Phosphorylation has also been identified as an antibiotic self-resistance mechanism^54^. A particularly relevant example of the latter is provided by the two kinases FomAB encoded in the BGC of the antibiotic fosfomycin that is widely used in human medicine (Supplementary Fig. 12). FomAB catalyze sequential phosphorylations on the phosphonic acid of fosfomycin^55–57^, which abolishes its bioactivity. But even a single phosphorylation was shown to be sufficient to inactivate fosfomycin, which left the role of the second phosphorylation somewhat surprising. The authors proposed that perhaps the second phosphorylation provides an intermediate with high free energy that is used to export fosfomycin diphosphate outside of the cell. Our work suggests a second potential explanation: the possibility that fosfomycin may be conjugated to another molecule after formation of fosfomycin-PPi analogous to the conjugation activation mechanism demonstrated here for fosfazinomycin. This hypothesis is supported by the identification of a previously uncharacterized FzmU homolog encoded immediately adjacent to the minimal fosfomycin BGC in *Streptomyces fradiae* (Supplementary Fig. 12, here termed FomU)^58^. FomU is conserved among putative *Streptomyces* fosfomycin producers, and may have escaped previous annotation for two reasons. First, determination of the minimal gene set required to produce fosfomycin showed that the gene encoding FomU is not essential^58^. Second, the DNA sequence that encodes FomU encodes another ORF (termed *orf2*^58^) on the opposite strand that appears to encode a putative dienelactone hydrolase (Supplementary Fig. 12). We did not find genes encoding homologs of the AS-like enzyme FzmA or conserved precursor peptides near these homologous fosfomycin BGCs. Thus, if the reactions of FomAB are to prepare fosfomycin for conjugation chemistry catalyzed by FomU, fosfomycin may not be attached to a peptide but to a different molecule.

Transient cytidylylation of phosphonic acids by nucleotidyl transferases has been reported for a few phosphonates. The cytidyl group then assists in substrate recognition by downstream enzymes before hydrolytic removal to arrive at the final natural product^59–63^. Cytidylylation has also been recognized as a potential activation mechanism toward conjugation of a phosphonate to hydroxy acids or carbohydrates (Supplementary Fig. 13)^64–66^, although the actual conjugation step has not yet been reconstituted. In comparison, the activation mechanism described here by FzmK and FzmF as well as the subsequent conjugation reaction catalyzed by FzmU is unprecedented. To the best of our knowledge FzmU represents the first example of a nucleotidyl transferase that utilizes a surrogate of NTP to perform NMPylation-like chemistry. While the RRE encoding gene (*fzmT*) is positioned right next to *fzmU* in all homologous *fzm* BGCs, our data suggests that the ligation activity of FzmU is not strictly RRE-dependent. Future experiments are necessary to determine whether FzmT improves the efficiency of FzmU-catalyzed conjugation.

Our attempt to identify dedicated peptidases for the fosfazinomycin pathway was not successful. Instead, the unique hydrazide linkage in fosfazinomycins enabled the utilization of general aminopeptidases to cleave off the leader peptide. *In vitro* liberation of fosfazinomycin B from modified FzmZ was achieved through sequential digestion with endoproteinase LysC and aminopeptidase I from *Streptomyces griseus*. We envision that a housekeeping aminopeptidase or endopeptidase/ aminopeptidase pair would be responsible for the production of fosfazinomycin B. Such a proteolytic step would need to be intracellular because the Val-tRNA-dependent enzyme FzmI is anticipated to convert foasfazinomycin B into fosfazinomycin A intracellularly.

The successful elucidation of the fosfazinomycin pathway was possible because of the genome mining efforts to uncover novel AS-like enzymes. Indeed, a plethora of AS-like enzyme-encoding BGCs that are predicted to be unrelated to lasso peptide production remain uncharacterized^30^. These include FzmA homologs encoded in BGCs that appear to also encode the phosphonate biosynthesis enzyme PepM (Extended Fig. 5) with downstream biosynthetic elements that are different from those present in the fosfazinomycin BGC. Functional characterization of these BGCs may further expand the chemical diversity of RiPP and phosphonate natural products.

## Conclusion

In this study, we established the final missing steps in the biosynthesis of fosfazinomycins, revealing how Nature makes the key acylphosphonohydrazide. Five enzymes construct the fosfazinomycin backbone by activation of the phosphonate to its pyrophosphoryl anhydride and conjugation of this intermediate by a nucleotidyl transferase to a peptidyl C-terminal hydrazide derived from a ribosomal scaffold peptide by an AS-like enzyme. This work expands the chemical space of reactions catalyzed by AS-like enzymes and nucleotidyl transferases, and establishes an unanticipated link between RiPPs and phosphonates, which may inspire the discovery of more RiPP-phosphonate hybrid natural products.

## Methods

### General

All reagents used for chemical synthesis and *in vitro* assays were purchased from Sigma Aldrich, Chem-Impex, Ambeed and Santa Cruz Biotech unless otherwise specified. NMR spectra were collected on a B500 Bruker Avance III HD or a B600 Bruker NEO NMR Spectrometer. MALDI-TOF MS analysis was performed on a Bruker UltrafleXtreme MALDI-TOF mass spectrometer.

### Bioinformatics

The SSN of the asparagine synthase protein family (PF00733) was generated with the EFI-EST webtool^32,67^. UniRef 50 (∼15,000 entries) was selected due to the large number of proteins in the family. The alignment score was set to 135 so that members with roughly over 40% sequence identity form clusters. The SSN was submitted to the EFI-GNT to access genome neighborhood information. The neighborhood information was set to 20 genes with co-occurrence cut-off set to 0%. Clusters with over 50% of members that colocalized with at least one known select RiPP biosynthesis protein (RRE, PF05402; LanB-N, PF04738; LanB-C, PF14028; LanC, PF05147; LanM, PF05147-PF13575; LanKC, PF00069-PF05147; MvdD, PF21068; YcaO, PF02624; MNIO, PF05114)^25^ were selected and manually inspected. To identify the precursor peptides of FzmA homologs, the RODEO webtool^33^ was used with customized configurations that extract ORFs with a length of 5-50 amino acids.

### Bacterial strains

*E. coli* DH5α was used for the construction and routine propagation of expression plasmids. *E. coli* BL21 (DE3) was used for heterologous expression of all proteins.

### Plasmid construction

Expression plasmids were constructed through Gibson assembly. All PCR amplifications were carried out with NEB Q5® High-Fidelity DNA Polymerase or Takara PrimeSTAR® Max DNA Polymerase Ver.2.

### Solid-phase synthesis of FzmZ peptides

Fmoc SPPS was performed using a Liberty Blue™ microwave peptide synthesizer (CEM). Fmoc-Arg(Pbf)-WANG resin (Aapptec) and hydrazine 2-chlorotrityl resin were used for the synthesis of FzmZ and FzmZ-NHNH_2_ peptide, respectively. The N-terminal methionine was omitted from both peptides in the synthesis sequence. Ethyl cyanohydroxyiminoacetate and *N*,*N’*-diisopropylcarbodiimide were used as coupling reagents. After the coupling sequence was completed, the synthesized peptides were cleaved from the resin with 2 mL of 95% trifluoroacetic acid (TFA), 2.5% H_2_O and 2.5% triisopropylsilane. After 3 h reaction at room temperature (rt), the supernatant was added to 20 mL of ice-cold diethyl ether. After centrifugation, the precipitated peptides were dried and redissolved in 2 mL of deionized water before HPLC purification.

### HPLC purification of the synthetic FzmZ peptides

The synthetic peptide samples were filtered with a 0.45 μm filter prior to HPLC purification with an Agilent 1260 Infinity III system on a Waters XBridge® Prep C18 column (particle size: 5 µm; dimensions 250 x 10 mm). Purification was performed at a flow rate of 2.5 mL/min using 0.1% formic acid in H_2_O as solvent A and 0.1% formic acid in acetonitrile as solvent B. The same purification method was used for the purification of both peptides: 0–5 min 5% B, 5–10 min 5–35% B, 10–40 min 35–45% B, 40–43 min 45–5% B. Fractions containing desired pure FzmZ peptides were confirmed by MALDI-TOF MS, pooled and lyophilized.

### Protein expression and purification

The expression plasmids for FzmA, FzmH, FzmT, FzmU, FzmK and FzmF were used to transform chemically competent *E*. *coli* BL21 (DE3) cells. Single colonies were selected to inoculate Luria-Bertani (LB) media containing 50 μg/mL of kanamycin (for pET28a vector) or 100 μg/mL of ampicillin (for pET15b vector), and the resulting media was incubated at 37 °C overnight with 220 rpm shaking. The overnight culture was then used to inoculate 3 L of LB media supplemented with the appropriate antibiotic and the inoculated medias were grown at 37 °C with 220 rpm shaking until the OD reached 0.6. Then the culture was chilled on ice for 30 min, induced with IPTG (final concentration 0.2 mM). The induced cultures were grown at 18 °C with 220 rpm shaking for 14-18 h. Then the cells were harvested with centrifugation (6,880 ×g, 20 min, 18 °C). The cell lysate was resuspended in buffer containing 50 mM Tris, 300 mM NaCl, 10 mM imidazole and 10% glycerol (pH 7.5) with the addition of 1 mg/mL of lysozyme and Pierce^TM^ Protease Inhibitor Mini Tablet. After resuspension, the cells were lysed by sonication, and the lysed culture was centrifuged at 31,240 ×g (45 min, 4 °C). The resulting supernatant was loaded onto a 2.5 mL Ni-nitrilotriacetic acid (Ni-NTA) resin gravity column. After the supernatant passed through, 10 column volumes (CV) of wash buffer containing 50 mM Tris, 300 mM NaCl, 30 mM imidazole and 10% glycerol (pH 7.5) were added to the column. Then the proteins of interest were eluted by the addition of 5 CV elution buffer containing 50 mM Tris, 300 mM NaCl, 300 mM imidazole and 10% glycerol (pH 7.5). The eluted proteins were concentrated to 2.5 mL with Amicon® Centrifugal ultrafiltration tubes, then buffer exchanged to buffer containing 50 mM Tris, 300 mM NaCl and 10% glycerol (pH 7.5). The concentration of the purified proteins was estimated with the nanodrop A280 protein quantitation method. The protein stock solutions were stored at -80 °C.

### Synthesis of Me-HPnA, Me-HPnA-PPi and fosfazinomycin B

The synthesis of Me-HPnA and fosfazinomycin B followed literature reported procedures^21,40^. The NMR data were consistent with reported values. To synthesize Me-HPnA-PPi^68^, 0.1 mmol Me-HPnA was dissolved in 30 mL of 100 mM triethylammonium bicarbonate (TEAB) buffer. The solution was concentrated under reduced pressure. After one more dissolving-concentrating cycle, the residue was dissolved in 5 mL of deionized water, lyophilized, and the residual water was removed by five times co-evaporation with acetonitrile. The resulting dry phosphonate triethylammonium salt was dissolved in anhydrous dimethylformamide (DMF), then 1,1’-carbonyldiimidazole (0.16 g, 1 mmol, 10 eq) was added. The reaction was stirred at rt for 1 h, then anhydrous methanol (0.08 mL, 2 mmol, 20 eq) was added to the reaction dropwise. After stirring for another 40 min, tributylammonium pyrophosphate (Ambeed; 0.25 g, 0.45 mmol, 4.5 eq) was added and the reaction was stirred for another hour. Then the reaction was quenched by the addition of 100 mL of deionized water, and the diluted mixture was loaded onto a 2.5 × 25 cm DEAE Sephadex A-25 column that was preequilibrated with 50 mM ammonium bicarbonate. The column was washed with 300 mL of 50 mM ammonium bicarbonate and 1100 mL of 100 mM ammonium bicarbonate, then Me-HPnA-PPi was eluted with a 1 L 200-650 mM gradient elution. Fractions containing desired product were lyophilized to give Me-HPnA-PPi in 18% yield. ^1^H NMR (500 MHz, D_2_O) δ 4.65 (d, *J* = 19.4 Hz, 1H), 3.73 (s, 3H); ^13^C NMR (126 MHz, D_2_O) δ 172.2, 69.8 (d, *J* = 151.3 Hz), 53.0; ^31^P NMR (202 MHz, D_2_O) δ 4.4 (d, *J* = 25.0 Hz), –5.7 (d, *J* = 16.0 Hz), –19.5 (dd, *J* = 24.8 Hz, 16.0 Hz).

### *In vitro* FzmA and FzmH assay

For FzmA assays, the FzmZ peptide (100 μM) was mixed with FzmA (50 μM) in reaction buffer (100 mM Tris, 150 mM NaCl, pH 8). The sample was supplemented with 10 mM glutamylhydrazine, 5 mM ATP and 20 mM MgCl_2_. For FzmH assays, the FzmZ-NHNH_2_ peptide was mixed with FzmH (100 μM) in reaction buffer (100 mM Tris, 150 mM NaCl, pH 8). The sample was supplemented with 1 mM SAM. For coupled FzmAH assays, both enzymes were mixed with FzmZ with the reaction component concentrations as listed above. The reactions were incubated at room temperature overnight before acid quench and LC-MS analysis.

### *In vitro* FzmF and FzmK assay

Me-HPnA (4 mM) was mixed with FzmK (20 μM) and FzmF (50 μM) in reaction buffer (50 mM sodium phosphate buffer, pH 8.0) in 500 μL scale. The reaction was supplemented with 12 mM ATP and 12 mM MgCl_2_ and allowed to incubate at rt for 5 h. Then the proteins were removed by 10 kDa Amicon® Centrifugal ultrafiltration tube, and the mixture was supplemented with 30 μL of *d*_6_-dimethyl sulfoxide before ^31^P NMR analysis.

### *In vitro* FzmFKTU cascade assay

FzmZ-NHNH_2_ peptide was mixed with FzmH (100 μM) in reaction buffer (100 mM Tris, 150 mM NaCl, pH 8). The sample was supplemented with 1 mM SAM. After 4 h incubation at rt, FzmF (50 μM), FzmK (50 μM), FzmT (100 μM) and FzmU (100 μM) were added with Me-HPnA (1 mM), ATP (2.5 mM) and MgCl_2_ (20 mM). The reaction was incubated at rt overnight before acidification to quench the reaction.

### *In vitro* peptidase assay for fosfazinomycin B production

The mixture of FzmZ-NHNHMe and FzmZ-fosB peptide (derived from the FzmHFKTU assays) was first digested with endoproteinase LysC (NEB) in a 1:50 (v/v) ratio. After overnight incubation at 37 °C, the mixture was first acidified to remove LysC. Then the pH of the solution was adjusted back to ∼ 8, and aminopeptidase I (A9934, Sigma) was added in a 1:10 (v/v) ratio with the supplementation of 1 mM CaCl_2_. After overnight incubation at 37 °C, the digestion was quenched by acidification prior to LC-MS analysis.

### LC-HRMS and MS/MS analysis of modified FzmZ peptides

An Agilent 1290 LC-MS QToF system was used for ESI-HRMS and MS/MS analysis. LC separation was achieved at 50 °C using a Phenomenex Aeris 2.6 μm PEPTIDE XB-C18 LC column at a flow rate of 1 mL/min using 0.1% formic acid in H_2_O (solvent A) and 0.1% formic acid in acetonitrile (solvent B) with the following gradient program: 0–2 min 5% B, 2–12 min 5–95% B, 12–15 min 95% B, 15–17 min 95–5% B. Mass spectra were collected in positive mode at 10 spectra/s and 100 ms/spectrum. Tandem-MS fragmentation was achieved at normalized collision energies of 20, 25 and 30. MS/MS analysis was performed using the Interactive Peptide Spectral Annotator^37^ and verified manually. For the analysis of fosfazinomycin B, which is very hydrophilic, LC separation was achieved at 30 °C using a Thermo Scientific™ Hypercarb™ Porous Graphitic Carbon HPLC column with the following gradient method: 0–4 min 5% B, 4–12 min 5–95% B, 12–17 min 95% B, 17–18 min 95–5% B.

## Data availability

The primary data for all figures are deposited in Mendeley Data at

## Acknowledgements

This manuscript is the result of funding in part by the National Institutes of Health (NIH; grant R37 GM058822) and the Howard Hughes Medical Institute. A Bruker UltrafleXtreme mass spectrometer used was purchased with support from the Roy J. Carver Charitable Trust (Grant No. 22−5622). W.A.v.d.D. is an Investigator of the Howard Hughes Medical Institute. The authors thank Yi Yang, Dr. Ryan Moreira and Dr. Songyi Xue for assistance with using the peptide synthesizer, and Dr. Lingyang Zhu for helpful discussions regarding NMR data.

## Author contributions

Experiments were designed by L.C., Z.H., C.P., K-K.A.W. and W.A.v.d.D. Experiments were performed by L.C., Z.H. and C.P. The manuscript was written by L.C. and W.A.v.d.D. The research was supervised by L.C. and W.A.v.d.D.

## Additional information

**Extended Data Fig. 1.**
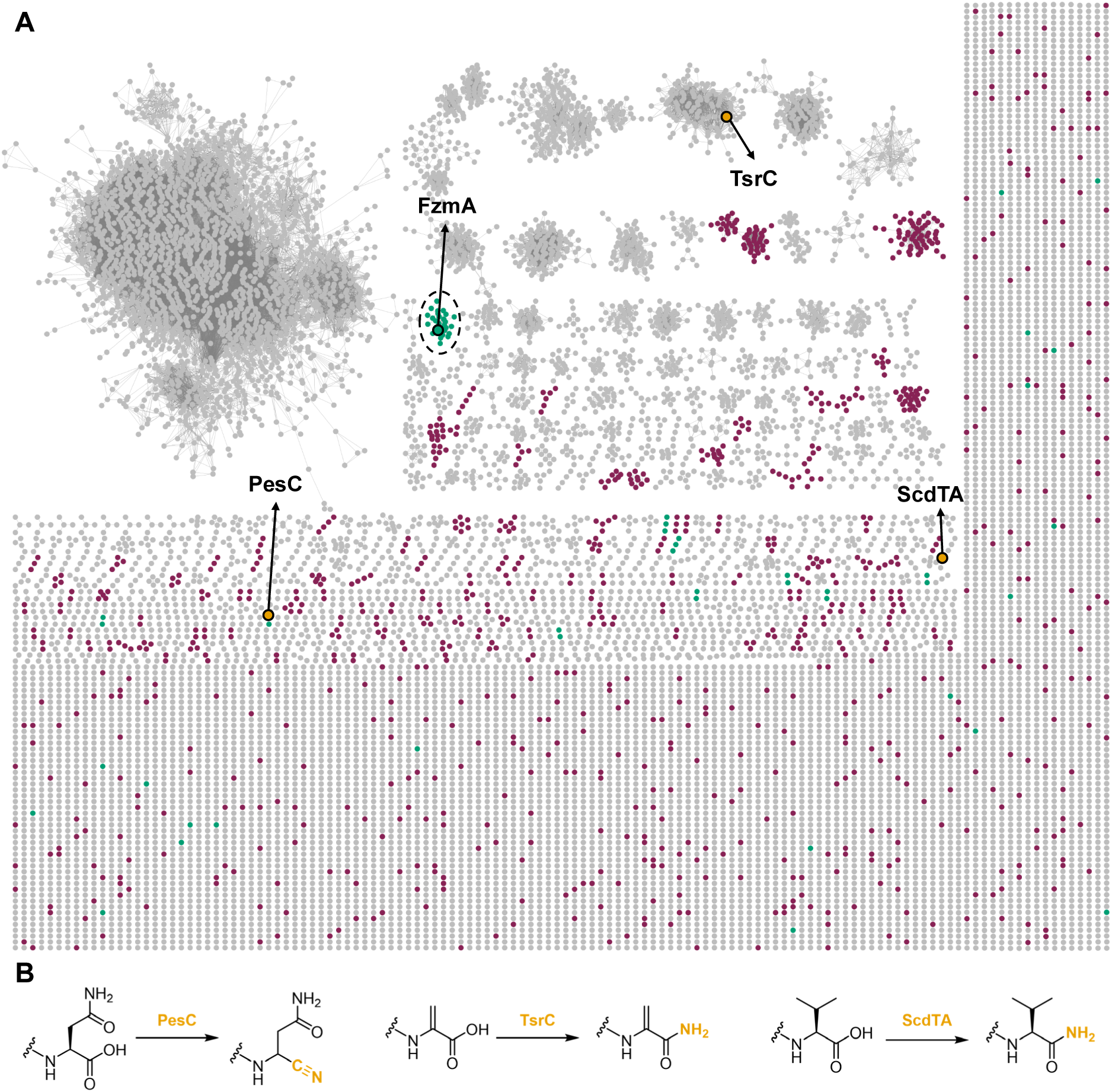
| AS-like enzymes in RiPP biosynthesis. **(A)** Sequence similarity network of the asparagine synthase (PF00733) family. The network was generated with the EFI-EST webtool^67^, covering ∼15,000 entries (UniRef 50) with the alignment score set to 135, resulting in members of > 40% identity to be arranged in clusters. Following genome neighborhood analysis, clusters with over 50% of members that colocalized with genes encoding at least one known RiPP biosynthesis element (see Experimental Procedures) are colored. Clusters related to lasso peptides are colored in plum, clusters that are predicted to be related to other types of RiPPs are in green. Three previously characterized enzymes that are not lasso cyclases are colored in yellow. The enzyme cluster that harbors FzmA is highlighted with a dashed circle (cluster 58). For a cytoscape file, see the Supporting Information of ref. 30. **(B)** The modifications catalyzed by PesC, TsrC and ScdTA (colored yellow in panel A). Only the C-terminus of the precursor peptide is shown.

**Extended Data Fig. 2.**
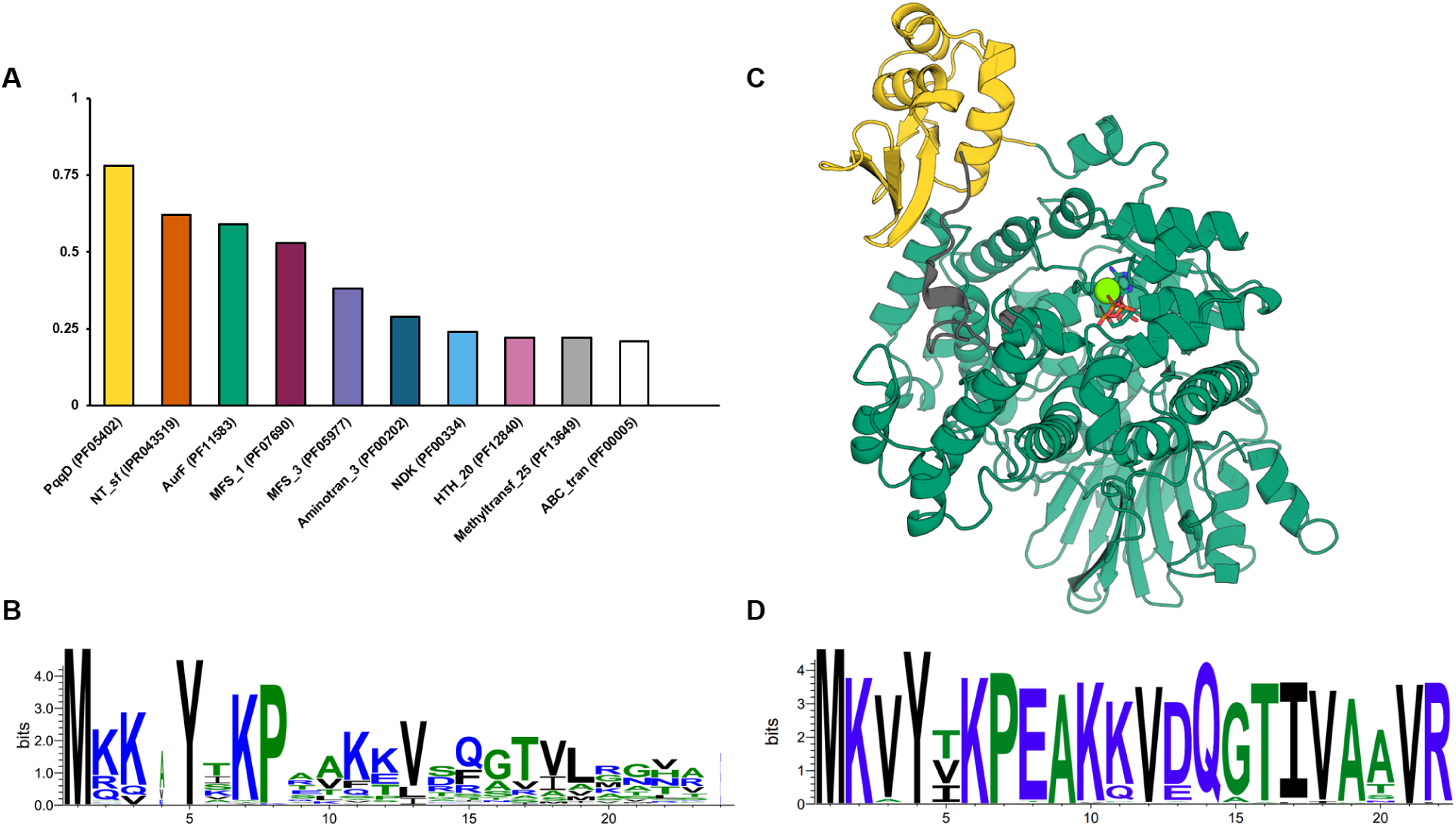
| Bioinformatic analysis of FzmA homologs. **(A)** Top ten protein families that are mostly frequently encoded in the genome neighborhoods of *fzmA* orthologs (89 members). **(B)** Sequence logo of short precursor peptides that are encoded in the genome neighborhood of FzmA homologs in the same cluster in the SSN showing a conserved YxxP motif. **(C)** The AlphaFold 3 predicted structural model of FzmA (green) bound with the precursor peptide FzmZ (gray), the C-terminal RRE domain of FzmA is colored in yellow. **(D)** Sequence logo^69^ of FzmZ-like precursor peptides from homologous *fzm* BGCs.

**Extended Fig. 3.**
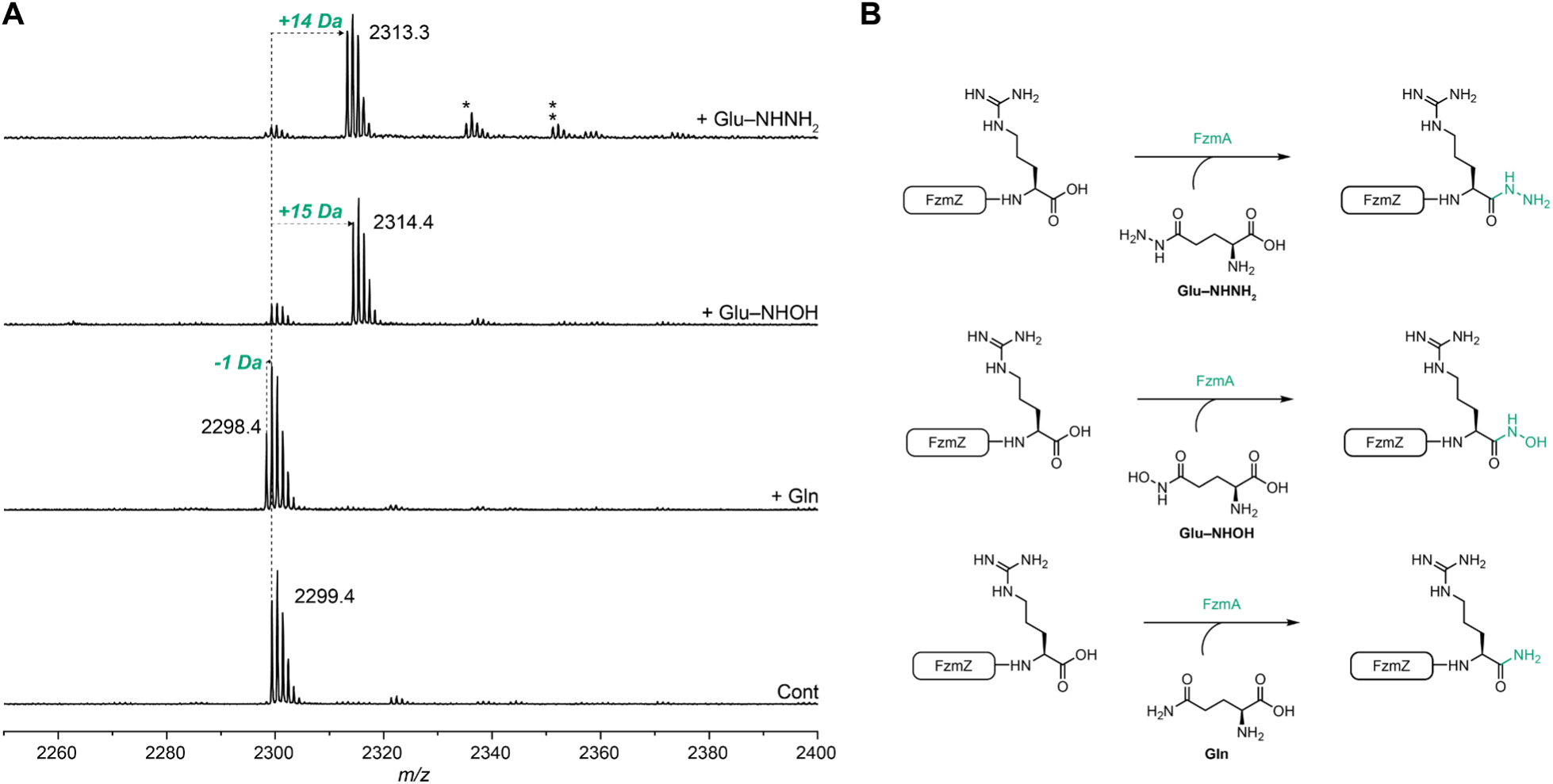
| The reactivity of FzmA with co-substrate analogs. **(A)** MALDI-TOF MS analysis of the *in vitro* FzmA reaction with FzmZ and L-glutamine or analogs of L-glutamine. Asterisk denotes [M+Na]^+^, double asterisk denotes [M+K]^+^. **(B)** Schematic representation of the reactions catalyzed by FzmA.

**Extended Fig. 4.**
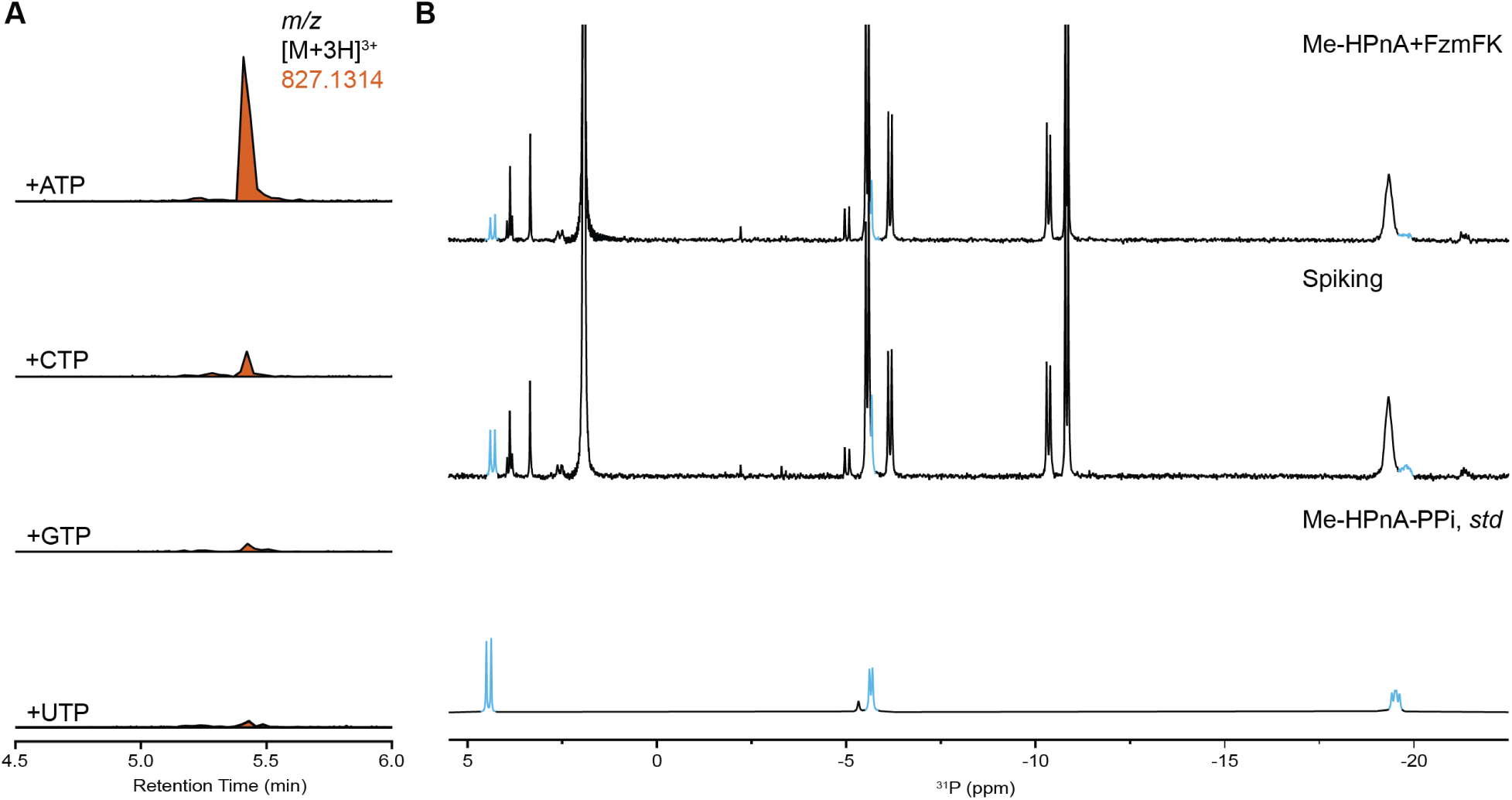
| Additional characterization of the Me-HPnA ligation. **(A)** LC-HRMS analysis of the *in vitro* FzmHFKTU reaction with FzmZ-NHNH_2_, Me-HPnA, and NTP (calculated mass for FzmZ-fosB [M+3H]^3+^: 827.1314). **(B)** ^31^P NMR analysis (taken in 6% *d*_6_-DMSO in H_2_O) of the *in vitro* FzmFK reaction with Me-HPnA as well as a spiking experiment with the synthetic Me-HPnA-PPi standard. The peaks of Me-HPnA-PPi are highlighted in blue.

**Extended Fig. 5.**
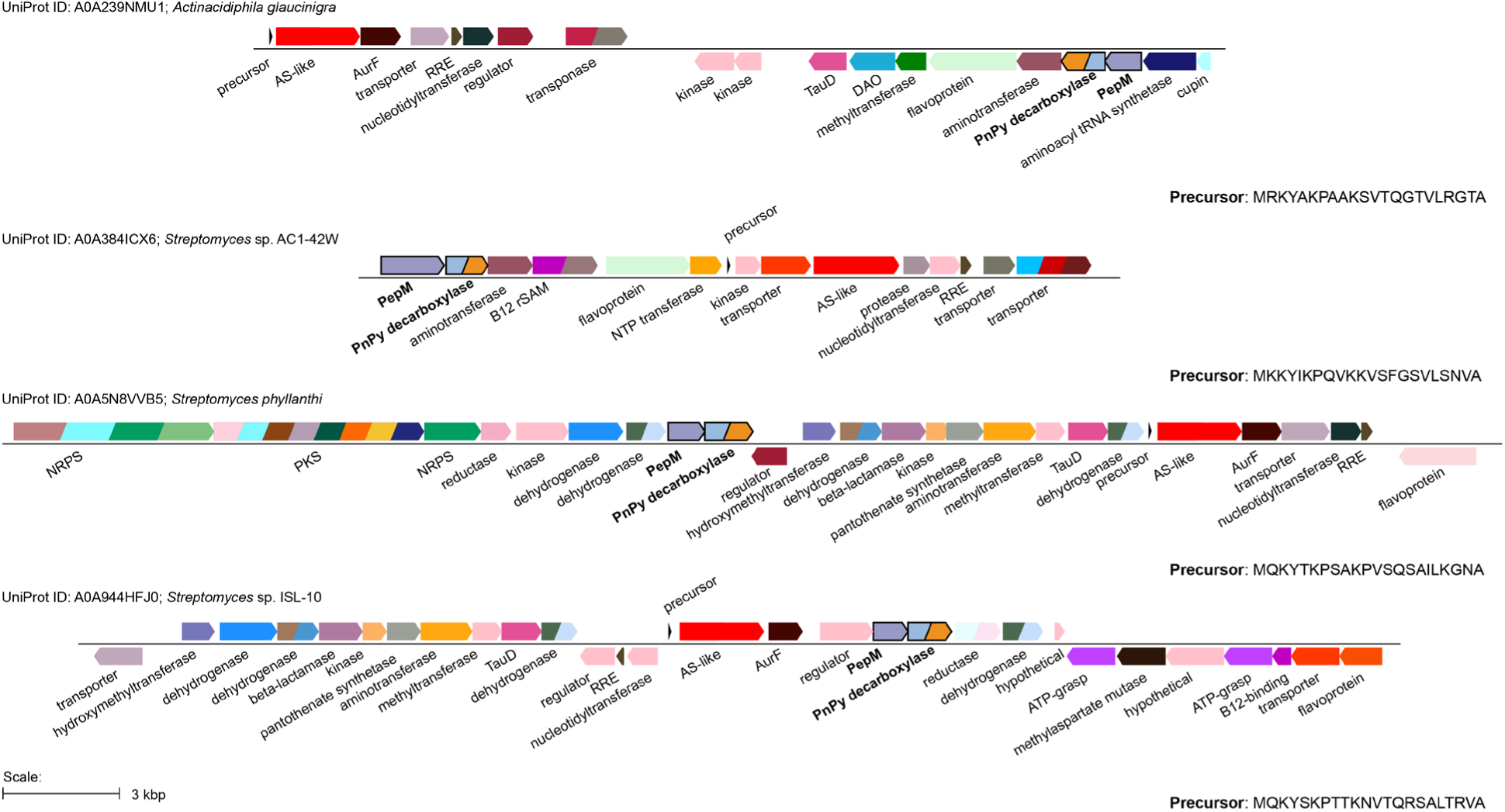
| Selected examples of genome neighborhoods of genes encoding FzmA homologs that harbor phosphonate biosynthesis machinery. The key enzymes that suggest phosphonate biosynthesis (PepM, PnPy decarboxylase) are highlighted in bold font. The Uniprot IDs of AS-like FzmA homologs as well as the hypothetical precursor sequences are listed for reference.

