## Supplementary Information for "Late-Stage Posttranslational Assembly of Fosfazinomycins"

### DNA constructs for expression of *fzm* BGC components in *E. coli*.

The DNA sequences encoding FzmH (WP\_053690973.1) and FzmK (WP\_159045979.1) were amplified from the genomic DNA of *Streptomyces* sp. WM6372 and NRRL F-2747. The DNA sequences encoding FzmA: WP\_030866196.1; FzmF: WP\_030866207.1; FzmT: WP\_053690966.1; and FzmU: WP\_147420027.1 were codon-optimized for expression in *E. coli*, synthesized, and inserted into vector pET-28a via Gibson assembly. This procedure yielded expression plasmids: pET-28a::<sup>C</sup>His<sub>6</sub>-FzmA, pET-28a::His<sub>6</sub>-FzmH, pET-28a::His<sub>6</sub>-FzmF, pET-28a::His<sub>6</sub>-FzmT, pET-28a::His<sub>6</sub>-FzmU and pET-15b::His<sub>6</sub>-FzmK. The gene sequences used in this study are shown below.

#### *fzmH*

```
atgacacggctcgtcgccgagctcgaacgactcggcctcgccgagttcctcgccgacttccccggggactcggtggacc
gccgccagtgggccgccggcgcccgcggcctcgacggtgagctgcgcgccctcgccgggttcctgctgctcggggagc
ccgcggcggccgaccggctgccgcccgccctcgccggggcgctgccctccctggtcaagagcggggtcgcgcaccag
cgcggggccgaggtccggctgatcgacgtcagcctgttccgctccgccggggtgtggctgttcgcggagccgccgagcc
ccttccccgtacgccactacttcggccgggactcgatcgcgctggcccggcgcaccggcttccgggcgggggagcgggt
cctcgacctgtgctccgggcctgggttccaggggctggtcgccgcacacggcggcgcccgggccacgctggtggagctg
ctgcccgagaccgccgaagtggcccggctcaacgcggagatcaacggggtggccgaccggatcgaggtcctcgtcgg
cgacctgtacgagccgctgccggccggggcgcgtccgtacgaccacgtcatcgcgaacatccccttcctgccgaccctc
gcgaccccgggcgaggaggccgcccacgagggcggcgaggacgggttcggcgtgggccgccgggtcctcgccggg
ctcccccggcacctgggcgcgggcggtacggcccatctgaccgcgctcctcctccagggggacgaacagctcgtcacg
gacggcgaactgcgggcctgggccgcggcgaacgggtacggcctgacggtcacgctgaccgagcggatggcggtgg
acgccgactccgacctcgtccagacgacggtgtcggaccacctggagaccgacccgggggccgaccccgaggagct
caccgccgccgccgtcgccgcgtacgcgagccgcggggcgacggccgcccgactggcctatctgcggatcgacgccg
ggatggccgactttcgtgtcctcgaccgcgcgggcgggtga
```

#### *fzmK*

```
atgatcgttgcgatggacggacccgacggcgcgggcaagagcacgcaggtcaagcgggtgcagagctgggccgagg
gccagggcctgtcgttccgcgtcgtcggcaagtggcaggtgttcgaggagtcccaggtgccgcaggcccgcttcctgcgc
ggtaccaccctcgacgaactgcgcgtctgcatcgccgagatgccgaacccggcccggatgctcttcctcggctggatgaa
caccctggccgccgagcgggcccgcgccgccgaggcggacctcgtcgtcctcgacggctactgggtcaagcacgcgg
ccaccgaactgctcgcgggctgtcccgaggagctggtggacgcgatcgtccacgccatggccccggtggacagcgtcg
tcttcttcgacgtcactcccgaggaggcactgcgccgcaagaagggcgacatcaccccgtacgaatgcgggcgggacc
cgcagtgccggccggagagcttcctgaagcaccaggccgccgtccgggacgtcatgctcgactgggcggcccggcgc
ggctgggacgtcgtacgggcgaacaccgccgaggcggcggcgcagcagctgcgcaaactgctggcgcccaagctgg
gcgtcgaggtcaacccccgccccacggtggccgacccgcgcaccgccgcggccgacccgcgcaccgccgtcgtcga
cgccccggcgggccactga
```

#### *fzmA*

atgtgcggaatagcaggatttgctcacactgacgggagcgcgctggctggttcagctgatgccatcttacgcgatatggcc
cacgccctgagaccacggggtccagatgatatgcagtttcatcatacaggcccagcgtgtatgtcctttacacgtctggcgtt
aattgatccggaaggtggcagacagcctttcattagcgaggacggcaatgtaattttagcggcaaatggggagatatata
actatcgcgaattgaagaaagggttcgaaggccgcgttcgctttcgctctgaatctgactgtgaagtgctgcttcacctgtata
tggaaaaagggctgagttttttggatgatgtacgtggaatgttcggtattgcggttatcgatctgcgggaacagcgccttttact
ggcgcgtgaccgcttaggtattaaaccgctttttattcatcgtaatcgcgatgcagttctgtttggttcggaggtgaaagccttat
ttgcccaccccgactgccctcgtgagctggattgggcgagagcacttgccgaccaaggcctttcatcagccccggttatga
gccacgacgaagcgatgacatggttcgtgggtgtagaccaggttcctgctggggccattgttgacattagcctgcgcgggg
gcgacacacgcgtccacagatactgggagttgccagctccggctgaagcaggtgggctgccggaatcgttttttgtcagat
caatgggagacctgctggctagcagcgttcgggaatgcttaatagcggatgccgagataggcgtctttctgtctggcggatt
ggactcggcagctataacaagtcttgcagcgcataccggtttgcacacatttagtgcattgactggtgccaccgttgtaaata
aagatgcgcaatacgcacgtgaaaccgctgacttacttgggctgccaaatcaccaagtcgcgttcggagcggaccgtatt
ccgagtcctgacgagtggcgtcgtctgctgtggcttatggaaacgccgctttgcggcccagagcagttttataaatccgaga
tgtatcgctttgcacgcgcggaaagacctgaactgaaagcgatccttttggggagcggtgctgatgaattttcagggggata
ttcagtagctctgagcggtgggggcgactgggaagactttgaaggaaatctgcgcacaatggcgcggcgtaaagctcttg
gttcacaatcggcgttaagcgtctggtgggaaggaacggaccttccgttgctgtccgatgacgcggtaggcgcatatgcgc
cgggcgcggtcgatgatccttatggctcatatctggcgtggaagttccgtgacatgcagcagtacaatttctgggtcgagga
ccgcactgcgtccggcaacggggttgaagctcgtgtgcctttcctggaccaccgtattattgaacttcttgcgacagttcctcg
tgcttatcggaaacggttattctgggataaacaagttatccgtgaagccgtacgtgatatccttccggccgaagtcatatcac
gtcccaaactggctttttatgagggcgaaggggtacgccatacgcatcgtacatttacaagaatgctggcgcagtcaggtg
atgaactggttgagcaagcattgagctcgcctagagcggcgcagtacctggatggtgcaaacatgcatcgtgcgctgcgg
agacttgaaagagatcagggcagcggtcatgtagaactgcttcttagagtcgtgaacctgggtattcttgatctgatggcggt
agatgtaccgggctctcgtgcccctaacggccctgctccgtacgaactgcgtgtaaccgactttgaaggcgcagaagttca
ggaactttttccgagacagcgtatcgaagaaacgagtgtgtttcgtctggcagaaggtgtgttggtgctggatgacgtaccc
ggagccaataccagctacgtagtcgttgatggtggtattcggttcgtgattgaccacctgacgcatgccgattggctgcggtt
acttagaggcttcgatggccgtcgccctttgtcagaggctttggatgccaccggatgcgccctggatgctatgagaccgctg
cttgatggctctttggaagctggtctgcttgaaagtgtggatgcacctgaaaaa

#### *fzmF*

atgcggcgggtcttgattgctaatagaggggccattgcagctcgcgcagcggatacttttcgcagcctgggatggagtccg
attgctgttgtcgcatcttcggacccgcagcgtctgcatgcgctggcagcggatggatgcgagtatctggagggcgagggt
ctggcacaaacatacgatcatgcaggtcgtatcgtggaggccgcgcggcgttgtggcgcagacgcagtgtacccgggct
atggtgcgttggctgaagatcccgaactgccagagctgcttgcagccgccggtattgcctttgtcgggccgaccgcagaa
gttttagccgccgctcgtgacaaagatcatgcggtcgctaccgcggatagacttggcctgccggttttaccccacgcgacc
gggcatgagcgcatcgctaaattagttgctgacatcggtctgccggttatcttaaaaccggtgactggttgcggaggtttagg
tgtgagaatcgcgcgcaccgaatctgaagcagaacgtgctcttgctttgattggtgctgacgaaggcggcggcgatgaag
gtggaggcggggaaggttggtatgcagaacggtatgttgaacatggcgccgttgtgggagttacggtagccgtagacgc
ggcagcaacagttcatccgctgggtgaacgggaatccctgcttgtggacggttcaatgaaattactggaagcaagtcccgt
gcgtggtgtggccccagaattggtagaacgcatgcgtgtcgacacagcccgtttggtgacaggtatgggtttgcgtggggtt

gcaaccgttgagtttatagtcggctcgggctccggtagtggatccggaagcgatgcaggcggtggagggcggcgtcact
ggttccttgaagtaaacggccgccttcccctggcatatcgcatgtgtgaggctcagaccggcctggacgttgttgcacttcag
atggagttagctagaggcgcacttccggcgccgggtcgcatccgtgtagatcgggatgcacattgtatggaagcgcgcttg
ttcattggaccagctgcgggtggagcctcaggtagaattacggctcttgaattagcaccggaacctggtgtcacttataattg
cgcgcttgatgttcgctctccggtggtgttcgataatattgtcactcaggtgttggcacttggcggaagccggggtgcagccg
ctggtgcagtgttacgggcagtagaaggttcaggtgtatctggtgttggtacgtgcgctgaggagattgcggtgtggctgcgt
ggcagcggtcttgtcccggagcggcttggctcccaggcattaggagtttaa

##### *fzmT*

atgccgctggatgaaccaccgctgggcaaagacgtgcaggttgccccgggctcagttgccagcctgccgctgaacgtga
aaatccgcaaccaccgcggcaccaccgtggttggcggctatgagcatttctttgagctgacggcctctgcggcgcatatct
ggcgccagattgacggcaaacgcacggtgagcgatatcgcggcgctgattgcccaggagtacgacatcgaccaggaa
agcgtggttcaggacatcgttgaactgtttaccgaactggcgcagcacgatgtggtgaacatcgatcagggtgacagccg
cggctga

##### *fzmU*

atgacttatgataccagtggtgtgattgctgaactgaaaagccgtaacctgctgccggatgcctatgaaacggtttttgccag
cggctcgctgatccgtggctggggcaacccgacaagcgatctggatgtgcatgtggtcacccgcgatgtgtggaccagc
accattgatgagaccaaccacgttgcgctggagccgaacaccctgcagtatgaaggcacctttgttgatggccgccgctg
ggatgttgaatactggacagccagccagattgaacaggtgctgagcaaagtgagccaggagcagtttgccgatggccg
cggcacctggcgcaccctgagctaccacgagattgcgctgctggagcgtctgccgtatgcggcggcggcagatgacggt
gcgtggctggaaggcacccgtaaacgcctggcggcgtcggcgcaccgcagcgtgctgattgtcaacagcctgaaacag
gcggacagctacaccgaagatgcggcgggtcagattgcccgcggtgacctgtacagcgcggtgattgccaccaagaa
cgcctttaaccactcggttgatgcgctgcaggccagcctgggccagttcggcagcctgtggccgaaatggcgcgcgcgtc
gcatggaaatcctggatccggatgtgctgccgttcgatgcctattgggcgattgaaacgatgcgcagctttgatccggacca
tccgcgcaaatgggttgaagaaacggtggcggtctgccagcgtattagcatggaagttagcgtttaa

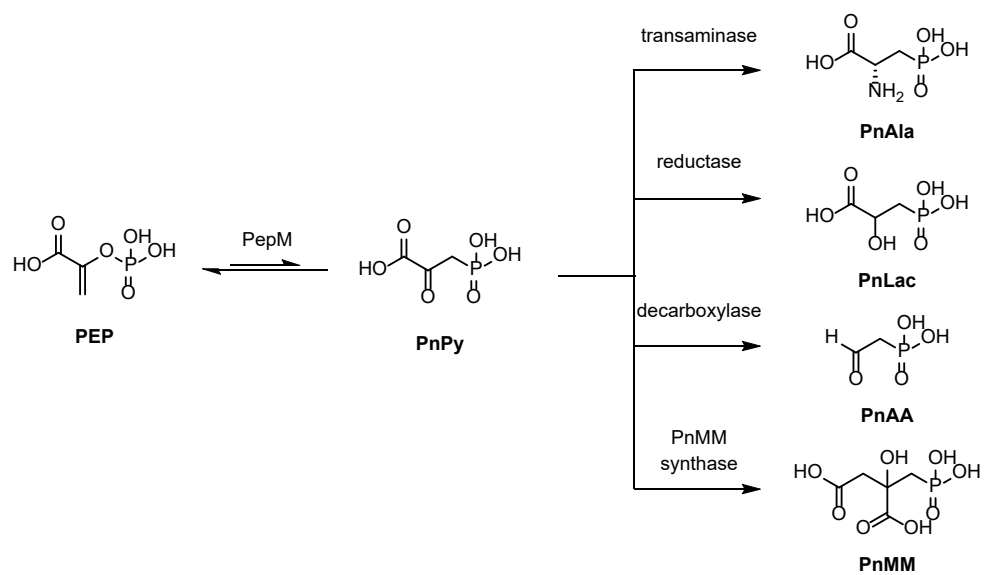

**Supplementary Figure 1.** The characterized early steps in the biosynthetic pathways of phosphonate natural products<sup>1</sup>.

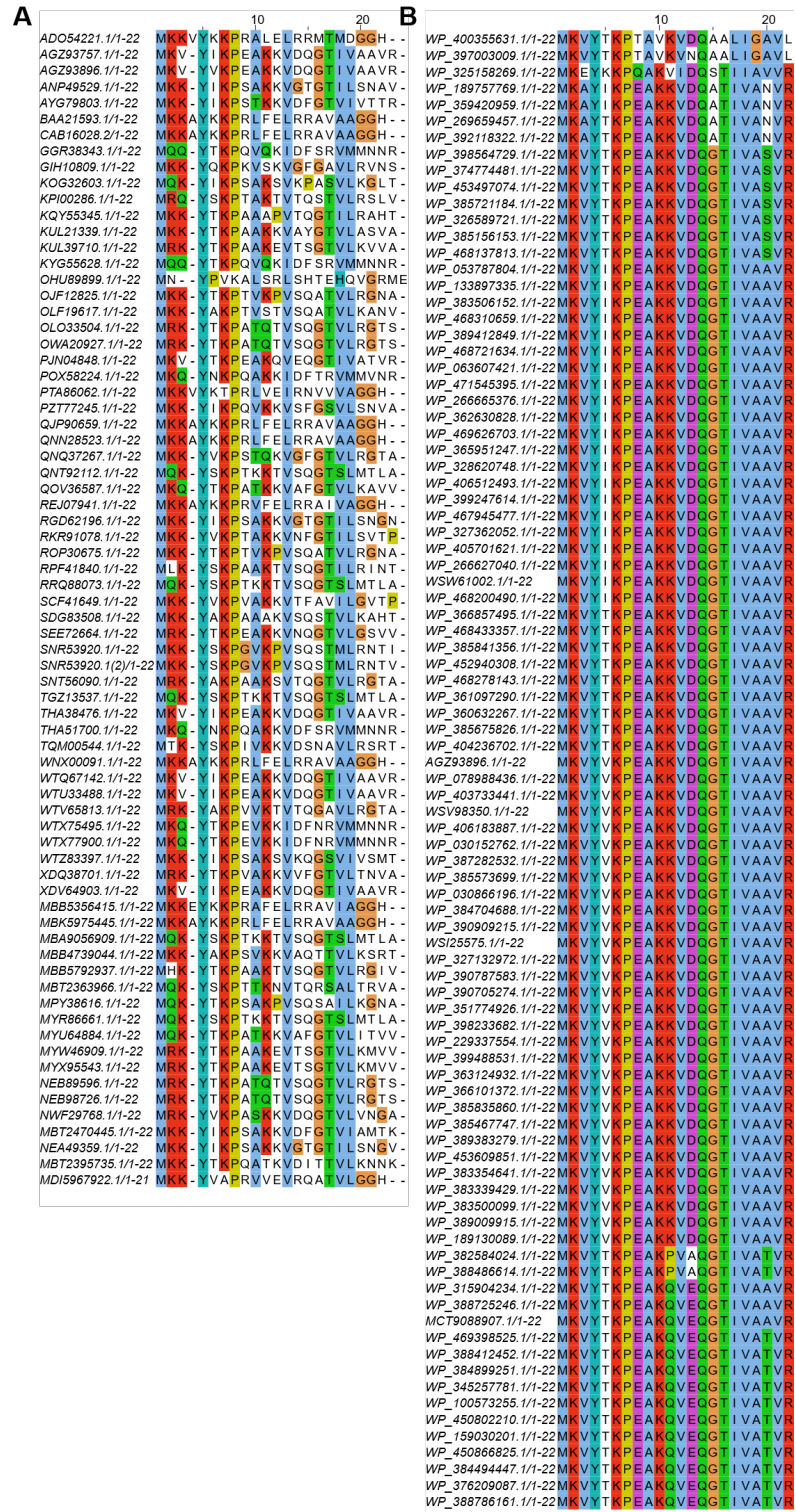

**Supplementary Figure 2. (A)** Multiple sequence alignment of the 72 precursor peptides identified from the genome neighborhood of AS-like enzymes from cluster 58 in the SSN. **(B)** Multiple sequence alignment of the 90 precursor peptides identified from homologous *fzm* BGCs. In both figures, the accession numbers of the neighboring AS-like enzymes are provided.

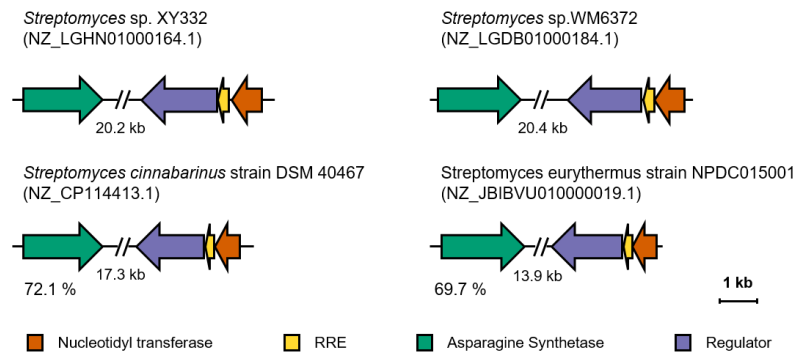

**Supplementary Figure 3.** Selected segments of the homologous *fzm* BGCs from two native fosfazinomycin producing strains *Streptomyces* sp. XY332 and *Streptomyces* sp. WM6372, and two other uncharacterized strains highlighting the conservation of a gene pair encoding an RRE and a nucleotidyl transferase. For the two uncharacterized strains, the identity of the FzmA homolog to FzmA from *Streptomyces* sp. NRRL F-2747 is provided for reference.

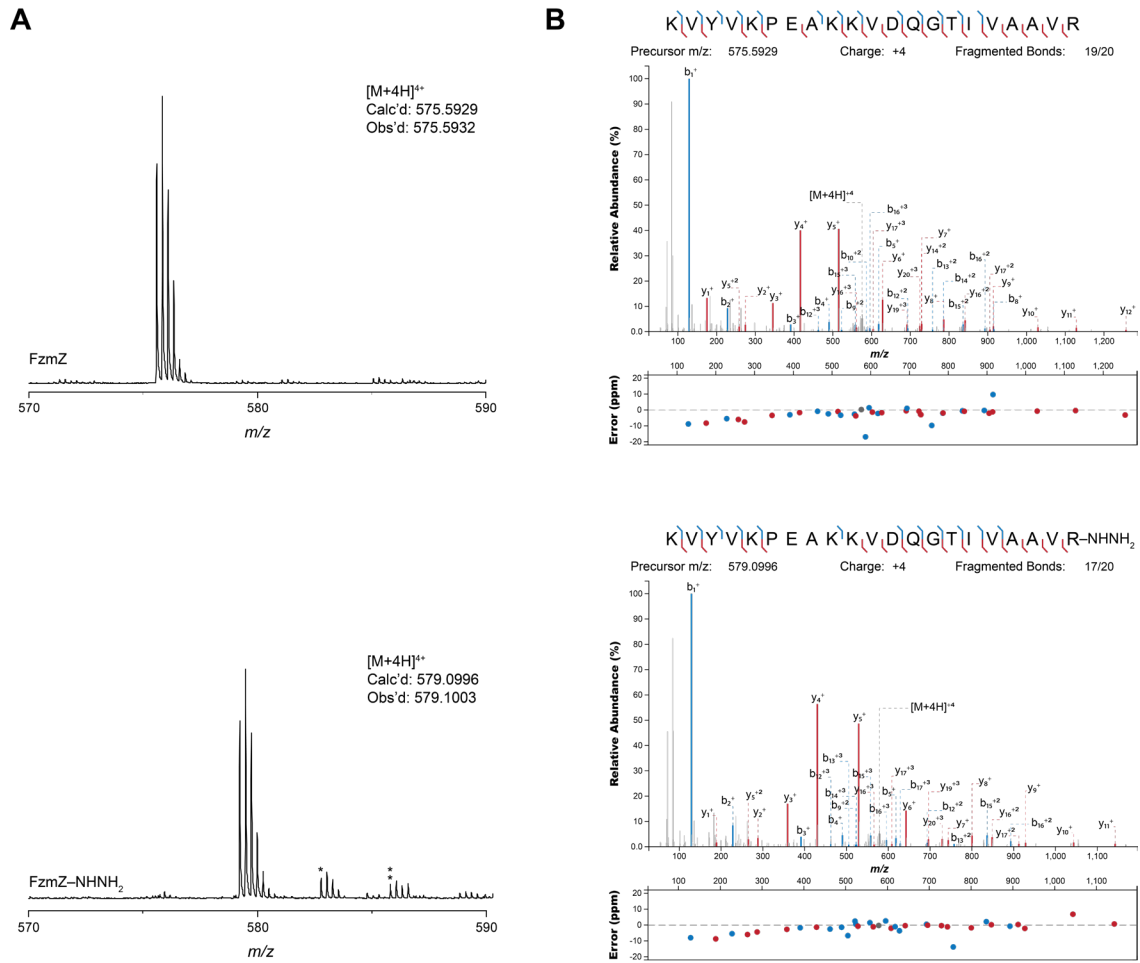

**Supplementary Figure 4. (A)** HRMS and **(B)** MS/MS analysis of chemically synthesized FzmZ (top) and FzmZ-NHNH<sub>2</sub> (bottom). Annotation of the tandem mass spectrum was performed using the interactive peptide spectral annotator<sup>2</sup>.

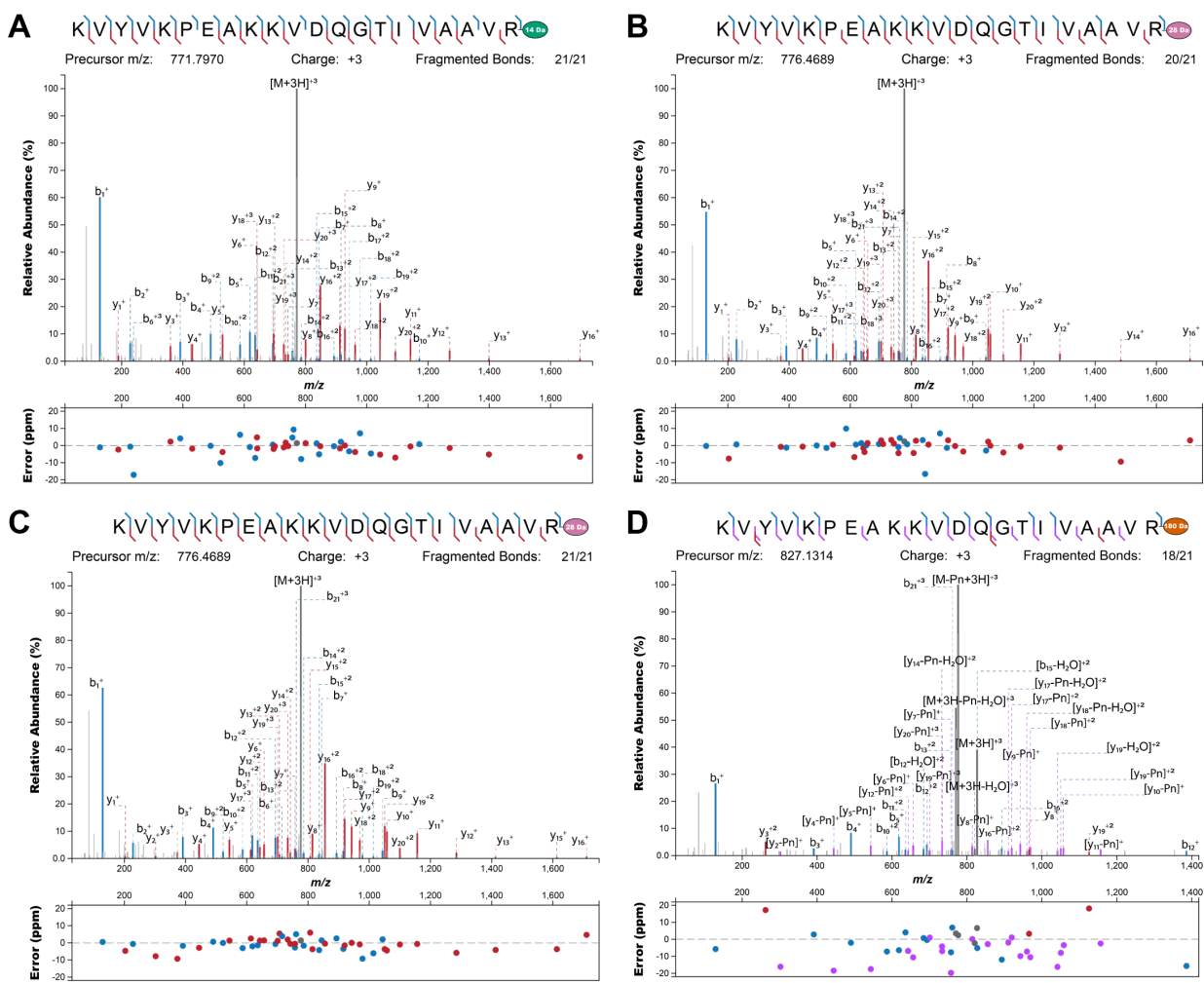

**Supplementary Figure 5.** MS/MS analysis of the products when FzmZ or FzmZ-NHNH<sub>2</sub> peptides were reacted with modifying enzymes: **(A)** FzmZ with FzmA; **(B)** FzmZ with FzmA and FzmH (FzmAH); **(C)** FzmZ-NHNH<sub>2</sub> with FzmH and **(D)** FzmZ-NHNH<sub>2</sub> with FzmHFKTU. Annotation of the tandem mass spectrum was performed using the interactive peptide spectral annotator<sup>2</sup>. The modifications introduced to the C-terminal carboxylate of the Arg residue are labelled as colored ellipses with the masses labelled. In Figure D, “-Pn” represents the loss of the Me-HPnA moiety (-152 Da).

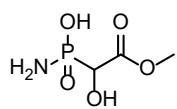

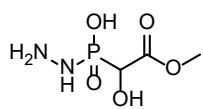

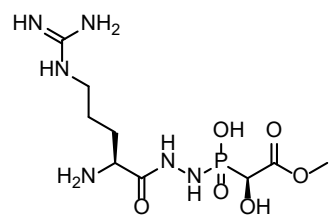

desmethyl-fosfazinomycin B

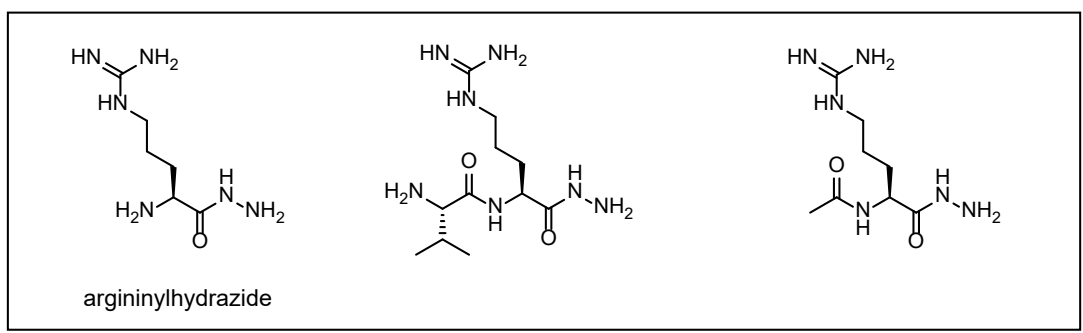

**Supplementary Figure 6.** The structures of substrate analogs that were previously used to test the methylation activities of FzmH<sup>3</sup>. Methylation on the terminal nitrogen was observed only when the three compounds highlighted in the box were incubated with FzmH and SAM.

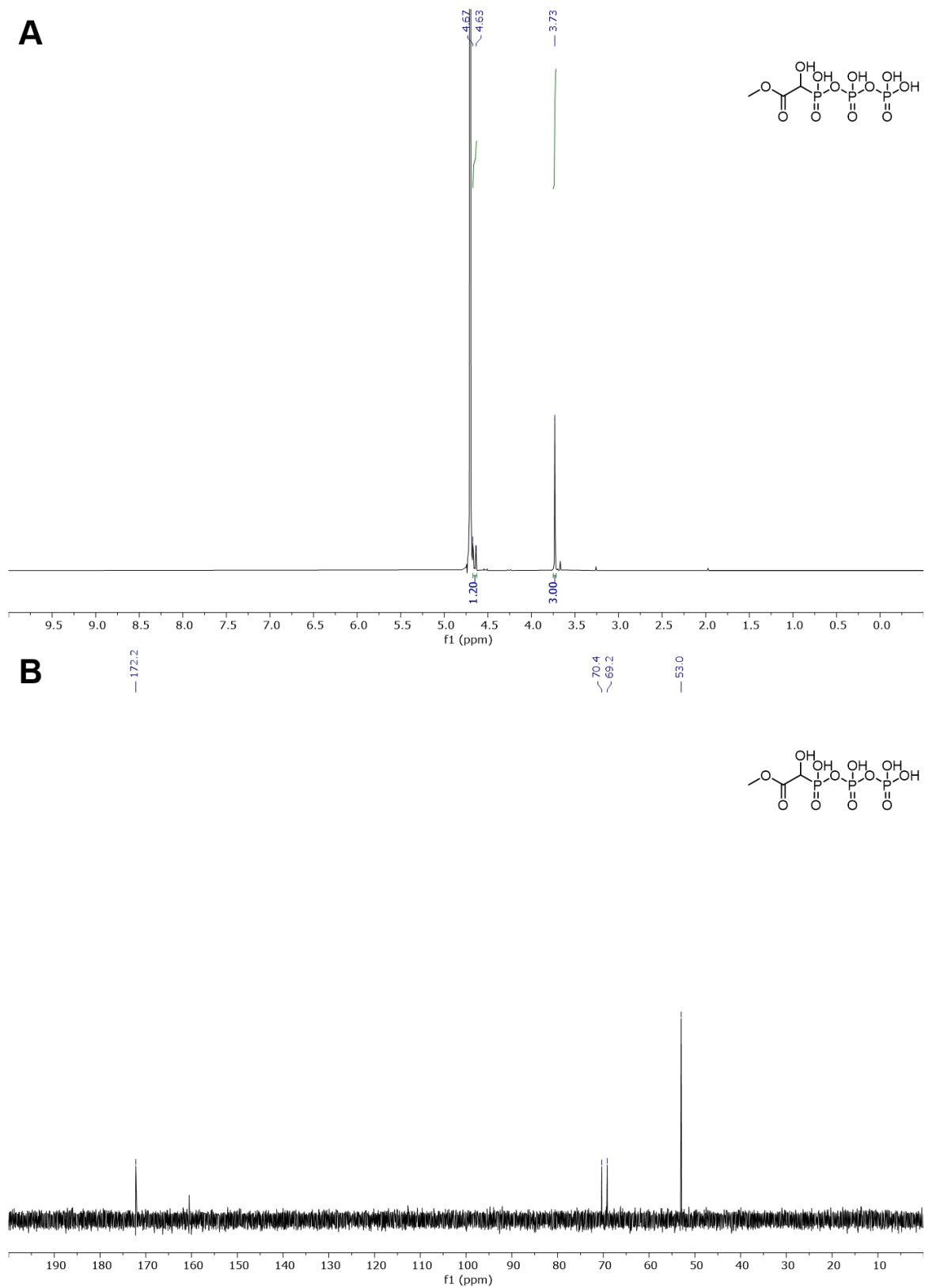

**Supplementary Figure 7. (A) $^1\text{H}$ and (B) $^{13}\text{C}$ NMR spectra (taken in $\text{D}_2\text{O}$) of Me-HPnA-PPI. In (A), the peak of the methine proton is very close to the water peak; see Supplementary Fig. 8B.**

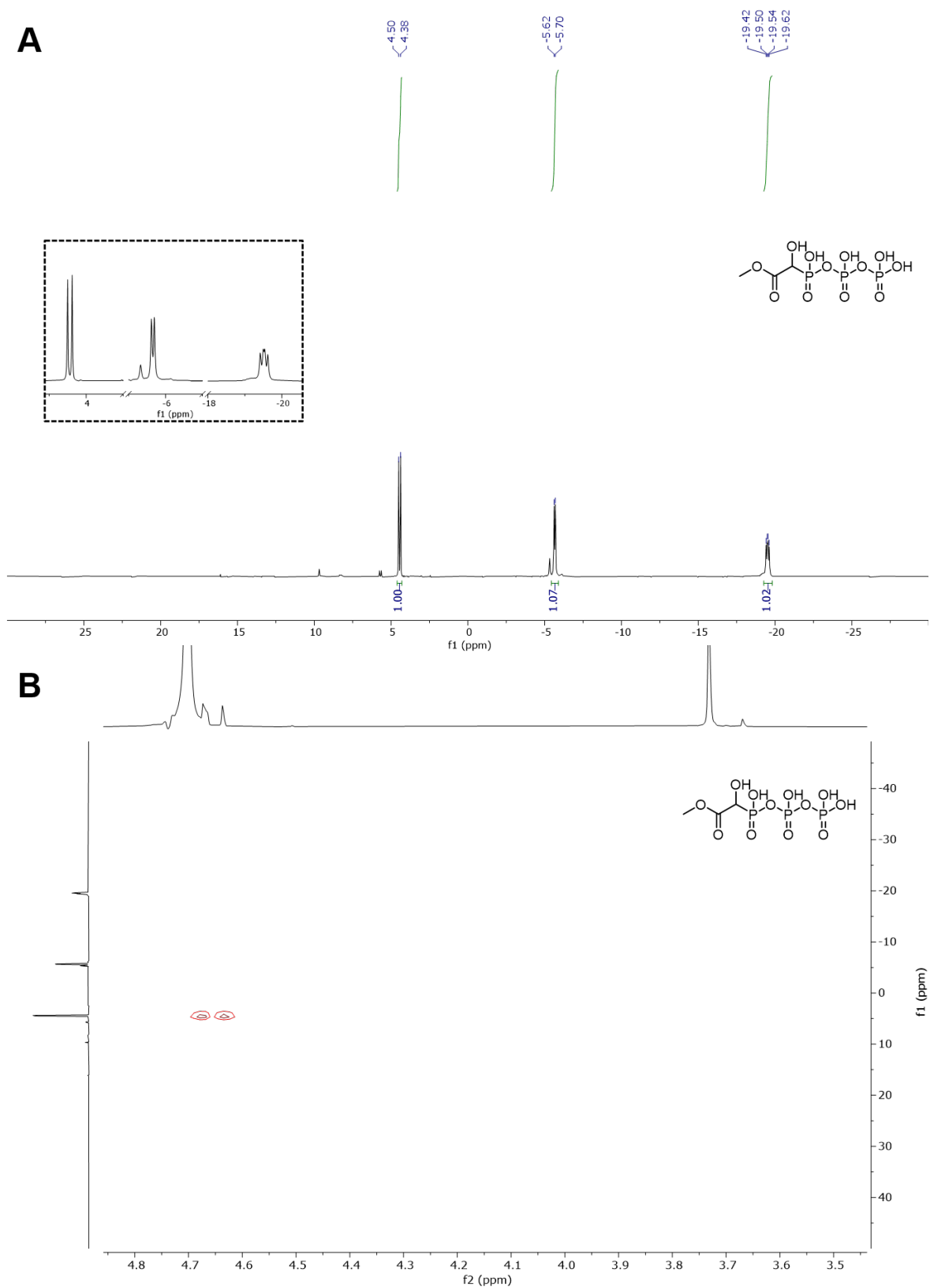

**Supplementary Figure 8. (A)** $^{31}\text{P}$ and **(B)** $^1\text{H}$-$^{31}\text{P}$ HMBC NMR spectra (taken in $\text{D}_2\text{O}$) of Me-HPnA-PPi.

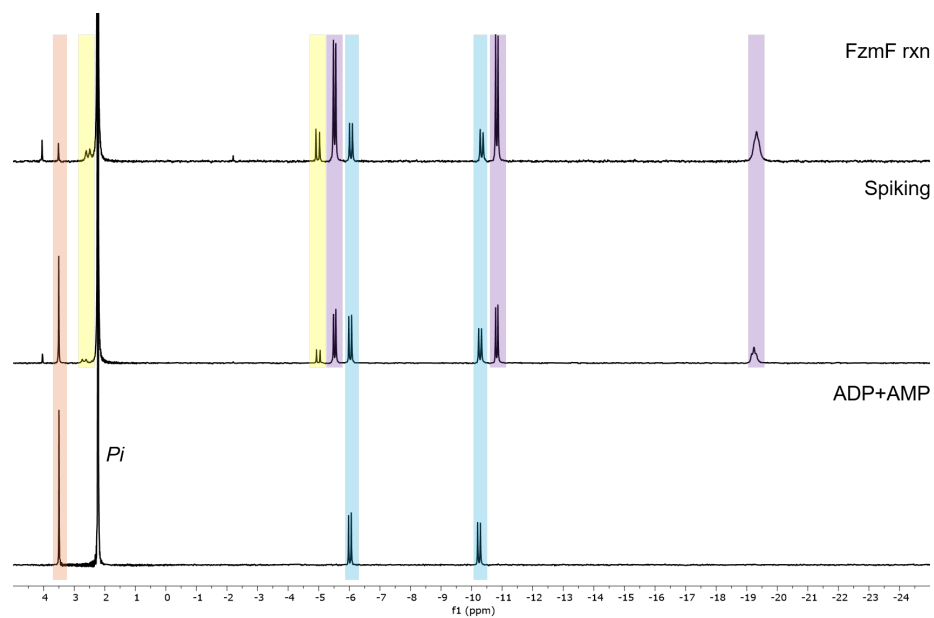

**Supplementary Figure 9. FzmF produces Me-HPnA-Pi.** $^{31}\text{P}$ NMR analysis (taken in $\sim 6\%$ $d_6$-DMSO in $\text{H}_2\text{O}$) of the FzmF reaction with Me-HPnA in the presence of ATP and $\text{Mg}^{2+}$. The reaction mixture was also spiked with ADP and AMP in NaPi buffer. The peak of AMP is highlighted in orange, the peaks of ADP are highlighted in blue, the peaks of ATP are highlighted in purple, and the peaks from the Me-HPnA-Pi product are highlighted in yellow.

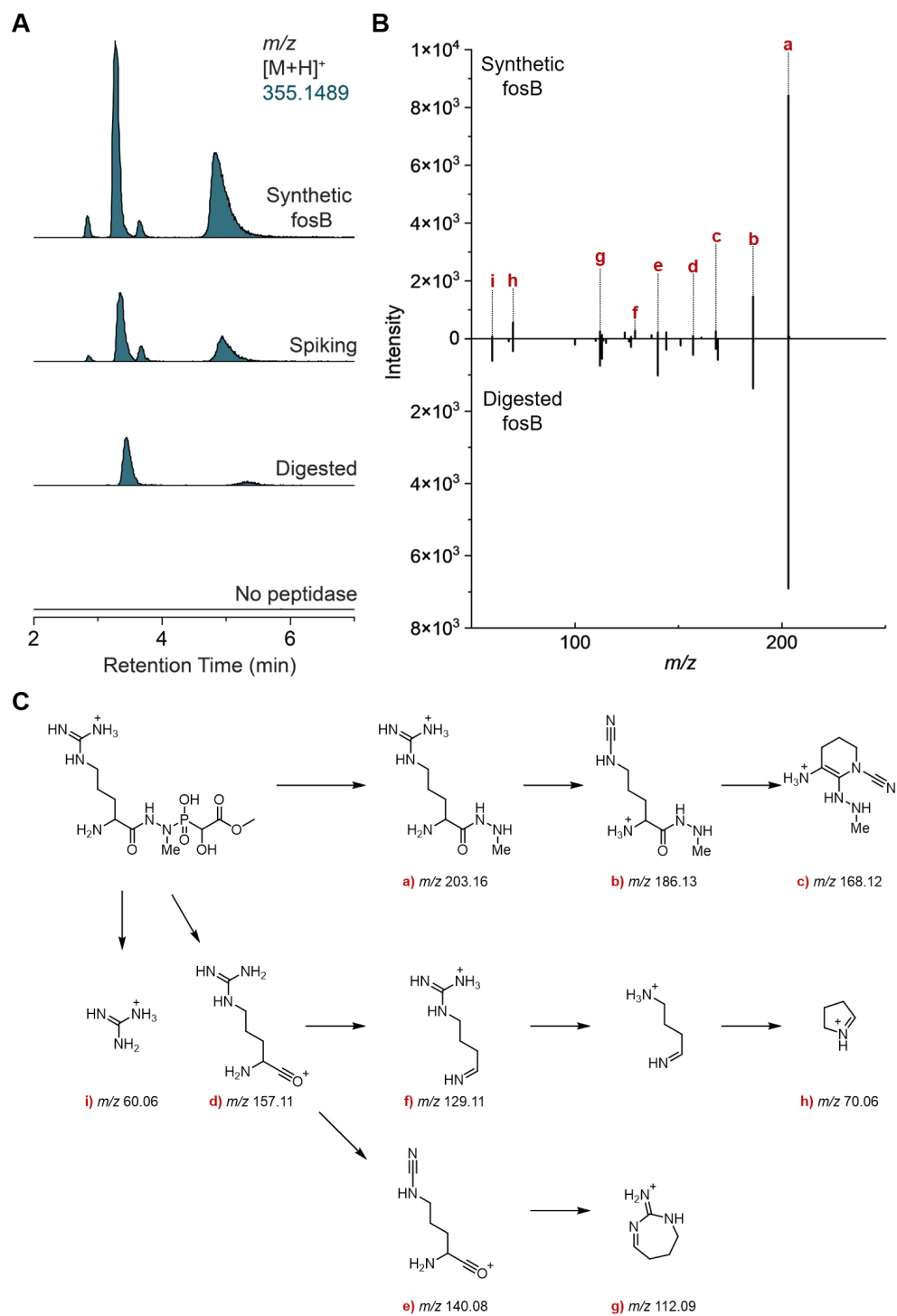

**Supplementary Figure 10. (A)** LC-HRMS analysis of synthetic fosfazinomycin B and epi-fosfazinomycin B (calculated mass: [M+H]<sup>+</sup> 355.1489) as well as fosfazinomycin B produced by LysC and aminopeptidase I digestion of the enzymatically prepared FzmZ-fosB peptide. **(B)** MS/MS analysis of the synthetic fosfazinomycin B standard (the isomer eluted at ~3.5 min) as well as fosfazinomycin B produced from digestion of the FzmZ-fosB peptide. **(C)** Proposed fragmentation pathway of fosfazinomycin B with assigned ions.

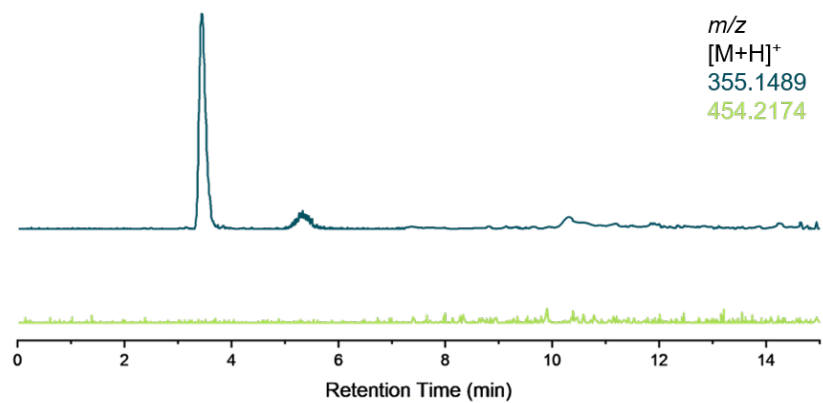

**Supplementary Figure 11.** LC-MS analysis of LysC and aminopeptidase I digestion of FzmZ-fosB peptide. Only the production of fosfazinomycin B (calculated mass: $[M+H]^+$ 355.1489) but not fosfazinomycin A (calculated mass: $[M+H]^+$ 454.2174) was observed.

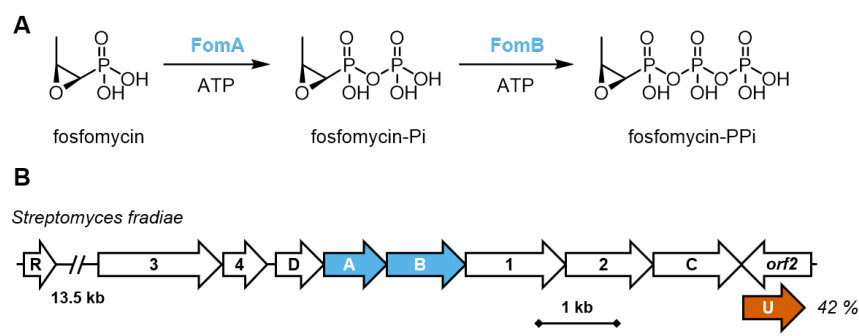

**Supplementary Figure 12. (A)** The sequential phosphorylations of fosfomycin catalyzed by FomAB. **(B)** The fosfomycin BGC from *Streptomyces fradiae*<sup>4</sup>. An uncharacterized homolog of FzmU (42 % identity), FomU, is encoded adjacent to the characterized fosfomycin BGC. In previous sequences of the fosfomycin BGC, an ORF on the opposite strand was annotated.

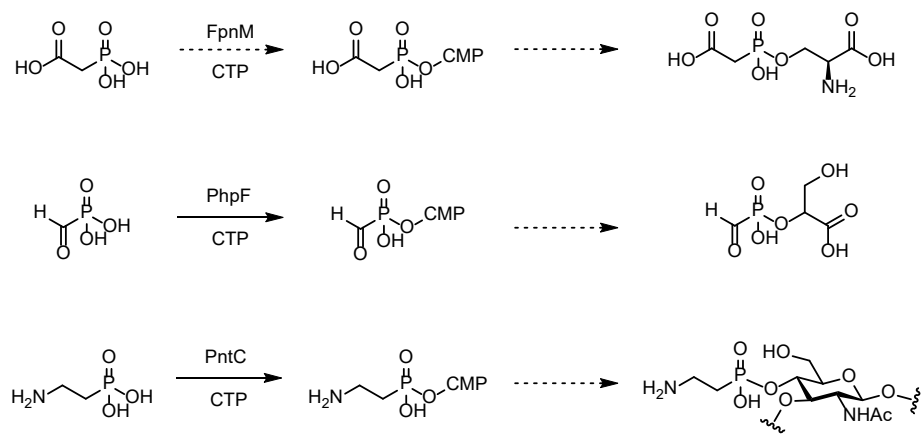

**Supplementary Figure 13.** Cytidylylation of phosphonic acid as an activation mechanism for phosphonate ligations during the biosynthesis of phosphonate natural products<sup>5-7</sup>.

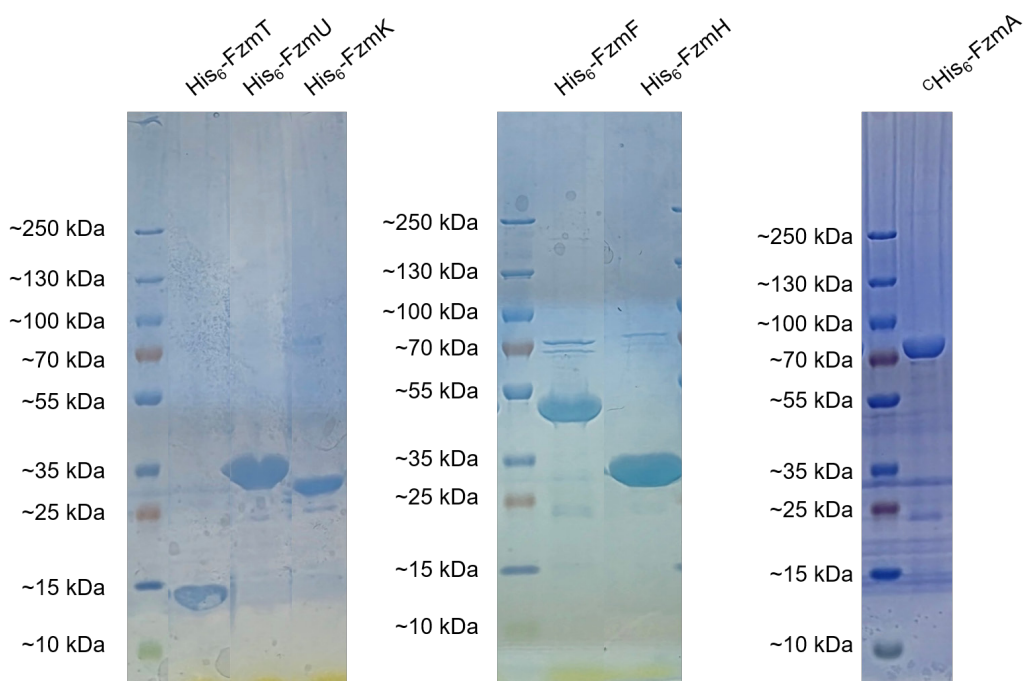

**Supplementary Figure 14.** SDS-PAGE of proteins used in this study: His<sub>6</sub>-FzmT (13.9 kDa), His<sub>6</sub>-FzmU (31.4 kDa), His<sub>6</sub>-FzmK (28.4 kDa), His<sub>6</sub>-FzmF (48.1 kDa), His<sub>6</sub>-FzmH (38.7 kDa) and C-His<sub>6</sub>-FzmA (82.4 kDa).
